# Defining hierarchical protein interaction networks from spectral analysis of bacterial proteomes

**DOI:** 10.1101/2021.09.28.462107

**Authors:** Mark A. Zaydman, Alexander Little, Fidel Haro, Valeryia Aksianiuk, William J. Buchser, Aaron DiAntonio, Jeffrey I. Gordon, Jeffrey Milbrandt, Arjun S. Raman

## Abstract

Cellular phenotypes emerge from a hierarchy of molecular interactions: proteins interact to form complexes, pathways, and phenotypes. We show that hierarchical networks of protein interactions can be extracted from the statistical pattern of proteome variation as measured across thousands of bacteria and that these hierarchies reflect the emergence of complex bacterial phenotypes. We describe the mathematics underlying our statistical approach and validate our results through gene-set enrichment analysis and comparison to existing experimentally-derived hierarchical databases. We demonstrate the biological utility of our unbiased hierarchical models by creating a model of motility in *Pseudomonas aeruginosa* and using it to discover a previously unappreciated genetic effector of twitch-based motility. Overall, our approach, SCALES (Spectral Correlation Analysis of Layered Evolutionary Signals), predicts hierarchies of protein interaction networks describing emergent biological function using only the statistical pattern of bacterial proteome variation.

## Introduction

A fundamental problem in biology is to understand how proteins interact to create a complex phenotype (Barabasi and Oltvai, 2004; Chuang *et al*., 2010; Hartwell *et al*., 1999; Costanzo *et al*., 2016). Biochemical and genetic studies have illustrated that complex behaviors emerge from a hierarchy of protein interactions: proteins interact to form complexes, complexes interact to form pathways, and pathways interact to create phenotypes (Papin *et al*., 2004; Ravasz 2009; Nurse 2008). Current strategies for identifying protein interactions span both experiment and computation. Experimental methods identify protein-protein interactions (PPIs) across different model systems and are continuing to evolve to be more high-throughput and comprehensive (Rajagopala *et al*., 2014; Schoenrock *et al*., 2017; Hauser *et al*., 2014; Koo *et al*., 2017; Luck *et al*., 2020). Computational methods leverage covariation between orthologous genes (orthologs) to predict PPIs (Eisen, 1998; Pellegrini *et al*, 1999; Enrich *et al*., 1999; Valencia and Pazos, 2002). Advances in computation as well as the breadth of available data have fueled continued innovation such as using statistical physical methods to infer PPIs from amino-acid coevolution (Croce *et al*., 2019; Cong *et al*., 2019; Green *et al*., 2021).

Pairwise PPIs derived from experimental and computational approaches are used to infer higher-order interaction networks (Szklarczyk *et al*., 2018). However, recent experimental work has shown that protein interaction networks defined only by binary interactions are incomplete, suggesting that important biological information lies in higher-order protein interactions (Kuzmin *et al*., 2018). Therefore, creating a model of genotype to phenotype relationships requires the ability to identify different ‘scales’ of interactions, from pairwise to higher-order, and relating these scales to describe the integration of pairwise interactions into higher-order interactions. We hypothesized that (i) pairwise and higher-order information could be directly extracted from the statistical pattern of covariation between orthologs and (ii) this information could then be used to create a single multi-scale hierarchical model describing the emergence of complex phenotypes from individual proteins.

We used Singular Value Decomposition (SVD) to spectrally analyze a large database of non-redundant bacterial proteomes and to define a set of components of ortholog covariation (an ‘SVD spectrum’). We found that covariation described by the SVD spectrum was distributed according to biological scale: top components were enriched for phylogenetic information, deeper components for higher-order protein interactions resembling pathways (‘indirect’ interactions), and deepest components for pairwise PPIs resembling physically interacting protein complexes (‘direct’ interactions). Second, we introduced the concept of ‘spectral correlations’, a metric representing the extent to which two bacteria or proteins share a similar statistical pattern of covariation within a specific region of the SVD spectrum. We found that machine-learning models trained on spectral correlation features could simultaneously predict indirect and direct PPIs with higher-fidelity relative to existing computational and experimental methods. Third, we introduced the concept of ‘spectral depth’—a way to relate spectral correlations between different positions in the SVD spectrum. Serially thresholding spectral depth defined hierarchically related protein interaction networks. We found our statistically derived hierarchies reflect the emergence of complex cellular phenotypes in bacteria as evidenced by gene-set enrichment analysis (GSEA) and comparison with experimentally derived hierarchies in the Kyoto Encyclopedia of Genes and Genomes (KEGG). The topology of these hierarchies were that bottom layers define protein interaction networks representing specific functions of protein complexes, middle layers define the integration of networks within bottom layers into broader functions resembling pathways, and top layers define the integration of networks in middle layers into high-level functions resembling phenotypes. Finally, we showed the utility of our approach by assigning global and local functions to an uncharacterized protein in *Pseudomonas aeruginosa* and validating these predictions experimentally. We call our approach for defining hierarchical protein interaction networks Spectral Correlation Analysis of Layered Evolutionary Signals (SCALES).

## Results

### Spectral decomposition of orthologous gene content among bacteria organizes ***covariation from phylogenetic relationships down to pairwise PPIs***

Orthologs are genes in different species that originated from a common ancestral gene and typically share a conserved core function. Assignment of orthologous gene groups (OGGs) is a robust and computationally tractable heuristic method for inferring orthologs and has been used extensively in comparative genomics (Overbeek *et al*.,1999). To sample variation in the OGG content of bacterial proteomes, we created a matrix, ***D^OGG^***, where each row is one of 7,047 UniProt bacterial reference proteomes, each column is one of 10,177 OGGs, and each entry is the number of times an OGG was observed in a proteome (**Figure 1A**, **Table S1**, **Figure 1 – figure supplement 1**, Materials and Methods) (The UniProt Consortium, 2019; Huerta-Cepas *et al*., 2017; Huerta-Cepas *et al*., 2019). We spectrally decomposed ***D^OGG^*** using SVD (Materials and Methods) (Klema and Laub, 1980) (**Figure 1 – figure supplement 2A-C**). SVD reveals patterns of correlations within the data by defining components of covariation and ordering them by their ability to explain the total data variance: SVD component 1 (SVD1) explains more data-variance than SVD2, SVD2 more than SVD3, etc (**Figure 1 – figure supplement 2D**). We observed that rows of ***D^OGG^*** corresponding to organisms sharing the highest level of phylogenetic classification, i.e. phylum, clustered together when projected onto SVD1, SVD2, SVD3, or SVD4 suggesting that the most dominant patterns of OGG covariation arise from global phylogenetic relationships (**Figure 1 – figure supplement 3**). The vast majority of the overall data variance (83%) was not explained by by SVD1, SVD2, SVD3, and SVD4 taken together. We next sought to systematically interrogate what biological information, if any, exists amongst deeper regions of the SVD spectrum of ***D^OGG^***.

**Figure 1.**
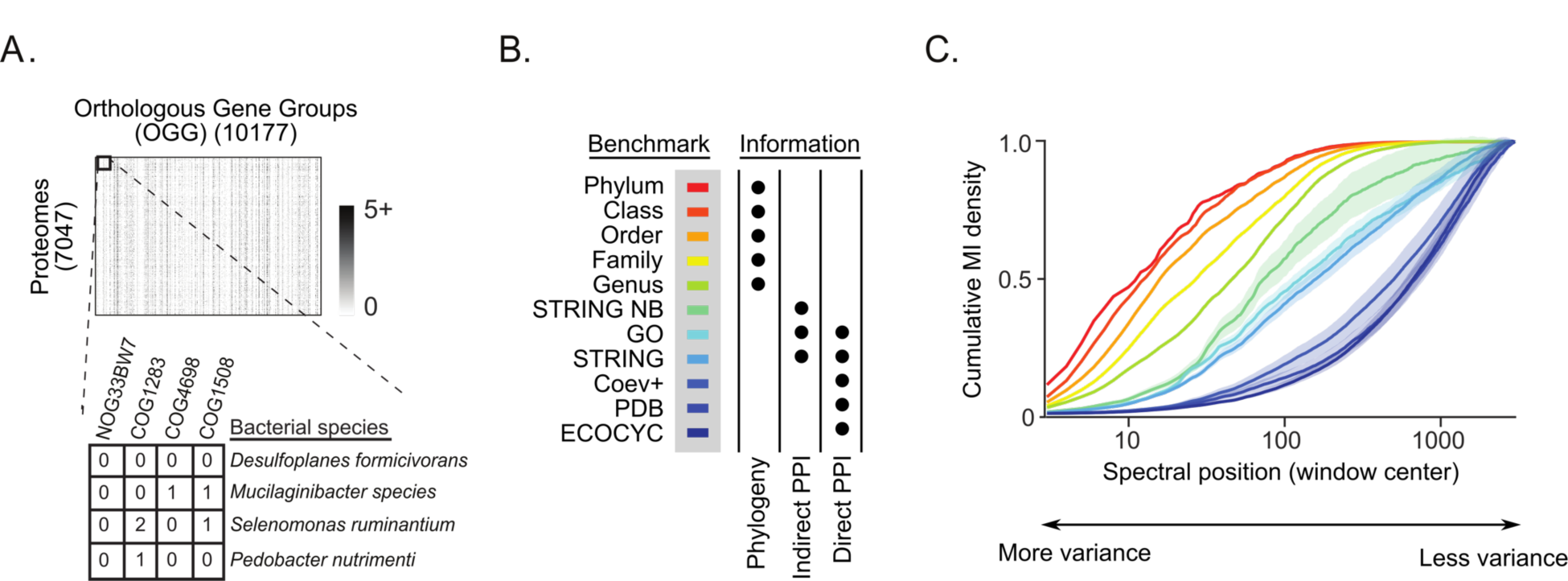
The SVD spectrum of OGG variation organized covariation according to biological scale. **(A)** The data matrix, ***D^OGG^***, contained the number of annotations of each of 10,177 orthologous gene groups (OGGs) within each of 7,047 UniProt bacterial reference proteomes. **(B)** Benchmarks were assembled to represent prior knowledge of phylogenetic relationships (Phylogeny), indirect PPIs (Indirect PPI), and direct PPIs (Direct PPI). For each benchmark, black circles indicate the types of information represented. **(C)** Cumulative distribution functions for the mutual information (MI cdfs) shared between the SVD components of ***D^OGG^*** and the benchmarks in panel B. Colors reflect the scheme in color legend of panel B. Shaded regions indicate ± 2 standard deviations surrounding the mean value for bootstraps of the benchmark. Components of covariation explain progressively less of the overall data variance with increasing spectral position.

To quantify biological information contained within different regions of the SVD spectrum, we computed the mutual information (MI) shared between known biological relationships spanning phylogeny to pairwise PPIs and the proximity between two proteomes or proteins as defined statistically by the SVD spectrum. A high MI value indicates that the statistical proximity reflects the known biological relationship, just as the clustering of proteomes on SVD1, SVD2, SVD3, and SVD4 reflected phylum classification (**Figure 1 – figure supplement 3**) In the following paragraphs we detail how we defined benchmarks of known biological relationships and statistical proximity between proteomes or proteins within a region of the SVD spectrum.

Benchmarks were assembled using existing biological databases to represent a hierarchy of organization from phylogenetic classification to indirect interactions in cellular pathways and to direct PPIs in protein complexes (**Figure 1B**, Materials and Methods). The NCBI taxonomy database was used to assemble five different phylogenetic benchmarks indicating if two bacteria share the same taxonomic substrings down to the levels of phylum, class, order, family or genus (**Table S2**) (NCBI Resource Coordinators, 2018). Pathway level benchmarks were assembled by mining indirect PPIs found in the STRING or GO databases (Szklarczyk *et al*., 2018; The Gene Ontology Consortium, 2020). Finally, protein complex benchmarks were assembled by incorporating direct PPIs identified in the Protein Databank (PDB), ECOCYC database, or by analyzing amino-acid level coevolution (Coev+) (Kesler *et al*., 2016; Cong *et al*., 2019). We focused our analyses on *E. coli* K12, a well-studied model organism for which a large number of high quality, experimentally supported PPIs are known (**Table S3**).

We quantified proximity within a specific region of the SVD spectrum of ***D^OGG^*** by introducing a metric we term ‘spectral correlation’ (Materials and Methods). In detail, application of SVD to ***D^OGG^*** produces two projection matrices ***U^OGG^*** and ***V^OGG^*** (**Figure 1 – supplemental figure 2A-C**). A row of ***U^OGG^*** contains the projections of a proteome onto each SVD component: column one onto SVD1, column two onto SVD2, and so on. Similarly, the rows of ***V^OGG^*** contain the projections of an OGG onto each SVD component. Spectral correlations are the Pearson correlations between two rows of ***U^OGG^*** or ***V^OGG^***. To compute spectral correlations for a specific region of the SVD spectrum, Pearson correlations are computed across the columns of ***U^OGG^*** or ***V^OGG^*** representing only the components of interest. The interpretation of a positive spectral correlation is that the two proteomes or OGGs are proximal when projected onto the specified set of SVD components. Because a single protein-coding gene can have multiple OGGs, we approximated the projection of a protein onto each SVD component by averaging the proteins’ constituent OGG projections and then computed protein-protein spectral correlations as above (**Figure 1 – figure supplement 4**).

**Figure 2.**
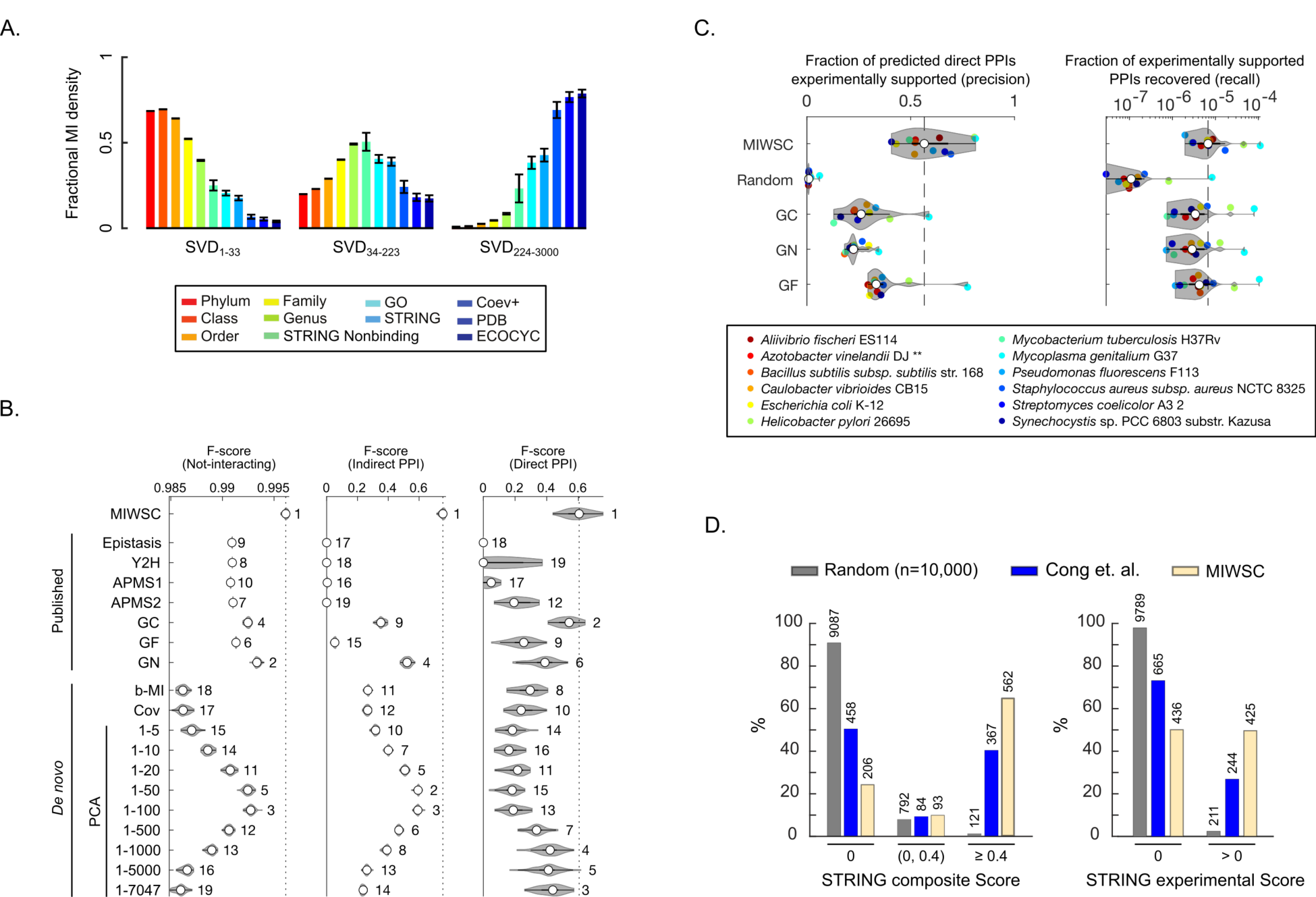
MI windowed spectral correlations (MIWSCs) enable accurate classification of protein pairs as either not-interacting, indirect PPI, or direct PPI. **(A)** Fractional MI density regarding each benchmark contained within spectral correlations computed across SVD1-33, SVD34-223, or SVD234-3000 of *D^OGG^*. Color scheme is defined in the legend and follows that of Figure 1B**,C**. **(B)** F-scores for predicting interaction classes for pairs in an independent validation set of *E. coli* K12 proteins using RF models trained on either MIWSCs, quantitative features of published datasets derived from experimental methods (gene epistasis [epistasis], yeast-two-hybrid [Y2H], affinity purification mass spectrometry [APMS1, APMS2]), quantitative features of published datasets derived from computational methods (gene cooccurrence [GC], gene fusion [GF], gene neighborhood [GN]), or quantitative features of established computational methods derived *de novo* from *D^OGG^* (binary mutual information [b-MI], covariation [Cov], Principle Components Analysis including components 1-k [PCA]). The violin plots describe the distribution of F-scores for models trained and validated on 50 random partitions of the gold-standard dataset (Figure 2 **– figure supplement 1**). Numbering indicates the rank of the median F-score for models trained on each feature (**Table S4**). (**C**) Precision (left) and recall (right) for direct PPI predictions in 12 phylogenetically diverse organisms using RF models trained on the MIWSCs of *E. coli* K12 proteins benchmarked against the experimentally supported PPIs in the STRING database. Comparisons are made to a set of 10,000 randomly selected pairs and to the ‘high confidence’ predictions in the STRING database subchannels for the methods of gene cooccurrence (GC), gene neighborhood (GN), and gene fusion (GF). Vertical dashed line indicates the median value for RF models trained on MIWSCs. ** in legend indicates an organism that was not part of the input dataset *D^OGG^* (**Table S4**). **(D)** Percent of predicted direct PPIs in *M. tuberculosis* H37Rv supported by an absent (0), low (0 to 0.4), or high (>0.4) composite score (left) or an absent (0) or present (>0) experimental subchannel score (right) in the STRING database. Comparisons were made between the methods of random selection (Random), amino acid coevolution (‘Cong *et al*.’, Cong *et al*., 2019), or RF models trained on MIWSC features of *E. coli* K12 proteins (MIWSC). Numbers of predicted interactions in each bin are indicated (**Table S4**).

We computed the MI shared between each benchmark and spectral correlations within all five-component windows of the SVD spectrum of ***D^OGG^*** (Materials and Methods). To estimate contributions of spurious correlations arising from finite sampling and overlap in OGG structure between proteins we computed the MI for a randomized projection matrix with bootstrap support (**Figure 1 – figure supplement 5**). We observed that the MI density (bits per window) for phylogenetic benchmarks declined rapidly as the spectral window was shifted deeper into the SVD spectrum, quickly converging upon the MI produced by spurious correlations (**Figure 1 – figure supplement 6A**). In contrast, the MI density decayed more slowly with spectral position for benchmarks of indirect PPIs and most slowly for benchmarks of direct PPIs. In fact, SVD2995- 3000 harbored significantly greater direct PPI MI than that produced by the model of spurious correlations despite accounting for only 0.021% of data-variance (p < 10^-243^ by pairwise Student’s T-test, **Figure 1 – figure supplement 6B**). These results suggest that the first 3000 components of the SVD spectrum of ***D^OGG^*** contain meaningful biological information.

We computed MI cumulative distribution functions (MI cdfs) for each benchmark across the top 3000 SVD components after subtracting the contributions of spurious correlations (Materials and Methods). Qualitatively, the MI cdf is the relative amount of MI gained as a function of depth; reaching a value of ‘1’ indicates that deeper regions hold no more biologically meaningful information regarding a benchmark. We observed that the MI cdfs for different types of benchmarks approached one in the following order: phylum, class, order, family, genus, indirect PPIs, mixed indirect/direct PPIs, and direct PPIs (**Figure 1C**). Of note, the MI cdfs for the three different benchmarks of direct PPIs—ECOCYC, PDB, and Coev+—were nearly superimposable, suggesting that the MI estimates were robust to benchmark source. Taken together, these results demonstrated that spectral decomposition of bacterial OGG variation using SVD organizes covariation from phylogenetic relationships down to pairwise PPIs.

### Using spectrally resolved covariation to train random forest models to predict indirect and direct PPIs in *E. coli* K12

A well-known challenge of using covariation to infer PPIs is that protein covariation can represent phylogenetic relationships, indirect interactions in pathways, direct interactions in protein complexes, or noise (Schafer and Strimmer, 2005; Sul *et al*., 2018; Nagy *et al*., 2020). The results above demonstrated that the SVD spectrum of ***D^OGG^*** separates covariation arising from these different sources. Therefore, we hypothesized that spectral correlations measured across specific SVD components could be used to accurately predict PPIs and assign them to a scale of organization, i.e. indirect vs direct PPI.

To test this hypothesis, we devised a multi-level classification task where a machine learning algorithm was challenged to classify pairs of *E. coli* K12 proteins as not-interacting, indirect PPI, or direct PPI (**Figure 2 – figure supplement 1**, Materials and Methods). For model training and validation, a gold-standard set of well characterized *E. coli* K12 protein pairs was assembled: 72,000 not-interacting pairs, 1,226 indirect PPIs, 72 direct PPIs. The indirect and direct PPIs were stringently chosen based on representation in multiple databases to reduce the rate of false positives in individual databases (Rajagopala et al., 2014). The not-interacting pairs were chosen by random selection. The relative numbers of not-interacting, indirect PPIs, and direct PPIs were chosen based on prior estimates of the true proportions of these interaction classes in biology (Rajagopala *et al*., 2014; Cong *et al*., 2019)The gold-standard dataset was randomly partitioned into training (60%) and validation (40%) sets. As will be described in detail below, spectral correlation features and various comparison features were extracted for the protein pairs in the gold-standard dataset. For each feature, Random Forest (RF) models were trained using labeled examples in the training dataset and validated using unlabeled examples from the validation dataset. Additional validation tasks performed included predicting out-of-bag examples in the training dataset and predicting examples in four additional comprehensive benchmarks (STRING Nonbinding: n = 14,793 indirect PPIs, GO: n = 79,794 indirect or direct PPIs, STRING: n = 14,793 indirect PPIs and n = 5,423 direct PPIs, PDB: n = 809 direct PPIs). RF model performance was quantified by computing F-scores for the predictions of each interaction class. F-score is the harmonic mean of precision and recall providing a holistic assessment of both the accuracy and completeness of each class prediction. Finally, the entire process of randomly partitioning the gold-standard dataset, training, and validation was repeated 50 times to assess the reproducibility of the resultant model performance.

To define spectral correlation features we computed spectral correlations across three sets of SVD components. The selection of these three sets was informed by the results of **Figure 1C**: SVD1-33 spanned up to the 25^th^ percentile point of the STRING nonbinding MI cdf, SVD34-223 spanned the 25^th^ to 75^th^ percentile points of the STRING nonbinding MI cdf, and SVD224-3000 spanned the 75^th^ percentile point to the point at which MI estimates converged upon that of the model of spurious correlations. SVD1-33, SVD34-223, SVD234-3000 isolated the majority of the MI related to either phylogeny, indirect PPIs, or direct PPIs respectively (**Figure 2A**). For each pair of proteins in the gold standard dataset, we computed spectral correlations across each of these three sets of SVD components (**Figure 2 – figure supplement 2A**). We term this set of three correlation values as the ‘MI windowed spectral correlations’ (MIWSCs) feature.

Comparison features were extracted from various existing datasets derived using established methods of PPI inference. These included the experimental methods of yeast-two-hybrid (‘Y2H’), affinity-purification mass spectrometry (APMS1’, ‘APMS2’) and gene epistasis (‘epistasis’), as well as the computational methods of phylogeny-aware gene co-occurrence (‘GC’), gene neighborhood (‘GN’), and gene fusion (‘GF’) (Szklarczyk *et al*., 2019; Rajagopala *et al*., 2014; Babu *et al*., 2014; Babu *et al*., 2018; Hu *et al*., 2009). As differences in the size and quality of the underlying dataset can influence the fidelity of computational PPI inference, we extracted additional comparison features directly from our dataset (***D^OGG^***). These additional features included binary MI (‘b-MI’), raw covariation (‘Cov’), and a Principal Components Analysis (PCA) based approach considering the top *k* SVD components (SVD1-k) (Materials and Methods) (Pellegrini *et al*., 1999; Franceschini *et al*., 2016).

In our multiscale classification task, the RF models trained on MIWSCs produced significantly greater F-scores across all three interaction classes compared to models trained on any of the other 18 different features in all validation tasks (**Figure 2B**, **Figure 2 – figure supplement 3**, statistical comparisons by Wilcoxon rank-sum test summarized in **Table S4**).

With regard to the PCA based approach, models trained on SVD1-5 performed at or below the median rank of all models across all three classes. F-scores for predicting indirect PPIs increased as components up to SVD100 were included (SVD1-100) without improvement of the prediction of direct PPIs. Including components beyond SVD100 increased the F-scores for predicting direct PPIs while decreasing F-scores for predicting indirect PPIs. Thus, the PCA based approach may not be ideal because models trained on the PCA-based features poorly predicted indirect PPIs, direct PPIs, or both. Taken together, these data show that the MIWSCs feature is the only one of the tested features that informs high fidelity PPI predictions in *E. coli* K12 across multiple scales of biological organization.

### RF models trained to predict PPIs in E. coli K12 using MIWSCs generalize across diverse ***bacteria***

To test generalizability of the RF models trained on MIWSCs in *E. coli* K12 to other organisms we predicted proteome-wide direct PPIs for 11 additional phylogenetically diverse bacteria, including one organism (*Azotobacter vinelandii*) that was not represented in ***D^OGG^*** (Materials and Methods). For comparison, we predicted direct PPIs for the same organisms using either random selection (‘Random’, n = 10,000) or by mining the high confidence interactions (confidence score > 0.7) in the STRING subchannels for the methods of GC, GN, or GF. To benchmark against orthogonal experimental evidence, we mined the experimental subchannel of the STRING database (Materials and Methods).

The median precision (5^th^-95^th^ percentile range in parenthesis) was significantly greater for direct PPIs predicted by the RF models trained on MIWSCs: 56.6% (41.0-81.2), 0.9% (0.6- 5.7), 26.1% (13.3-56.9), 22.4% (18.2-34.1), or 33.6% (29.5-74.6) for MIWSC RF models, random selection, GC, GN, or GF respectively (**Figure 2C**, left panel, one-sided Wilcoxon rank sum test with multiple comparison found in **Table S4**). In addition, the median recall was significantly greater for RF models trained on MIWSCs (**Figure 2C**, right panel, **Table S4**). The recall values were low across all methods, which may reflect the previously reported high number of false-positives in experimental databases (Rajagopala *et al*., 2014; Cong *et al*., 2019). Nevertheless, the MIWSC RF models predicted a median (IQR in parenthesis) of 897 (551 to 1609) direct PPIs per proteome.

In addition, we performed a head-to-head comparison for predicting direct PPIs in *Mycobacterium tuberculosis* H37Rv using the MIWSC RF models versus the method of Cong and colleagues that infers direct PPIs from proteome-wide amino acid coevolution (Materials and Methods) (Cong *et al*., 2019). We found that the MIWSC RF models exhibited a significantly greater precision and recall when benchmarked against the STRING composite score, as done previously by Cong and colleagues, or against the STRING experimental scores (**Figure 2D**, Chi-squared test found in **Table S4**).

Taken together, these results suggest that the RF models trained to predict *E. coli* K12 PPIs using MIWSCs were robust and generalizable across diverse bacteria.

### Spectral decomposition of protein domain content organizes covariation according to ***biological scale and informs accurate indirect and direct PPI predictions***

We sought to test whether the results we observed above were specific to choosing OGGs as a feature of orthology. Protein domains are conserved parts of proteins that have been previously used as a feature of bacterial orthology (Mistry and Finn, 2007). We defined a new matrix, ***D^domain^***, where each row is one of the 7,047 proteomes used in the ***D^OGG^*** matrix, each column is one of 7,245 unique protein domains, and each entry is the number of times a domain appears in a proteome (**Figure 3A**, **Table S5**).

**Figure 3.**
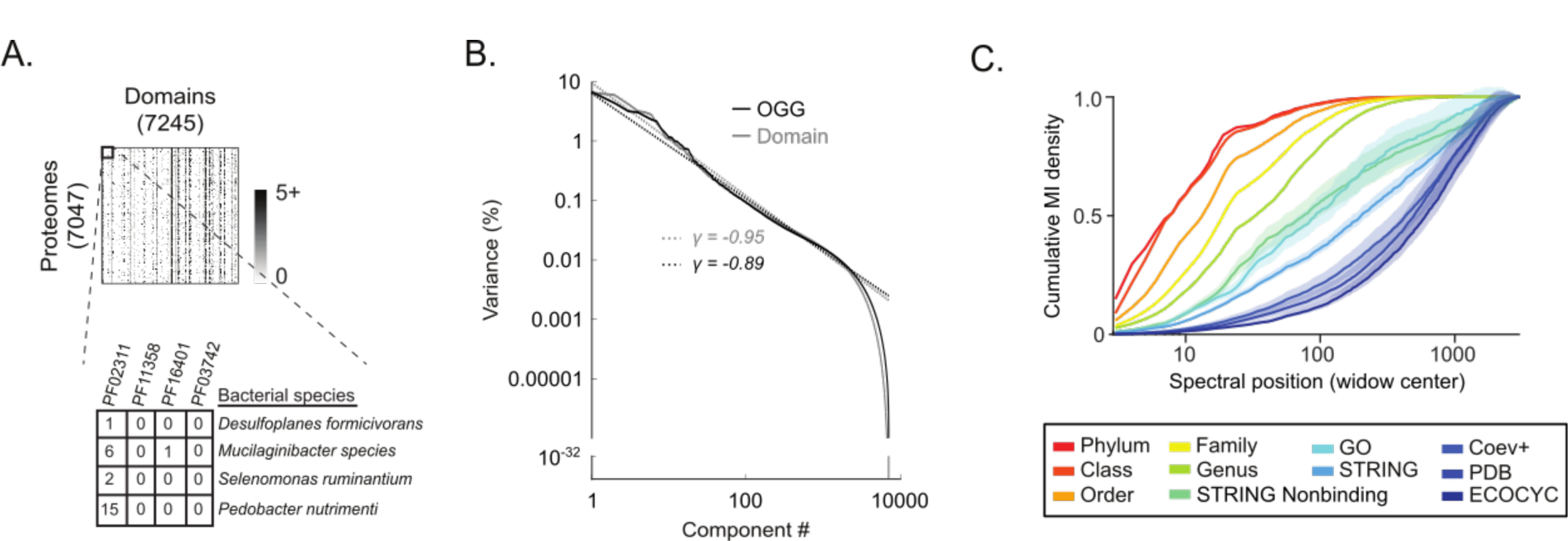
The SVD spectrum of protein domain variation organized covariation according to biological scale. **(A)** The domain content matrix, ***D^domain^***, contained the number of annotations of each of 7,245 conserved protein domains within each of the 7,047 UniProt bacterial reference proteomes. **(B)** Percent variance explained per component versus component number for the SVD spectra of ***D^OGG^*** (black) and ***D^domain^*** (gray). Dotted lines represent a power-law distribution with the indicated exponent (*γ*). **(C)** Cumulative distribution functions for the mutual information (MI cdfs) shared between the SVD components of ***D^domain^*** and the indicated benchmarks. Shaded regions indicate ± 2 standard deviations surrounding the mean value for bootstraps of each benchmark (STAR Methods).

The SVD spectrum derived from ***D^domain^*** was nearly superimposable on that of ***D^OGG^***, suggesting that the statistical structure of covariation is similar across these different orthologous features (**Figure 3B**, Materials and Methods). Similar to our analysis of ***D^OGG^*** described above, we quantified the MI shared between the various benchmarks of prior biological knowledge and spectral correlations within all 5 component windows of the SVD spectrum of ***D^domain^***. Again, we observed that the MI density (bits per window) for phylogenetic benchmarks declined rapidly as the spectral window was shifted deeper into the SVD spectrum (**Figure 3 – figure supplement 1A**). In contrast, the MI for indirect PPI benchmarks declined more slowly and the MI for direct PPI benchmarks remained statistically significant until at least SVD3000 (Paired Student’s T-test see **Figure 3 – figure supplement 1A,B**). MI cdfs were computed for each benchmark and found to mirror those derived for ***D^OGG^***: ordered according to phylum, class, order, family, genus, indirect PPI, mixed indirect/direct PPI, direct PPI (**Figure 3C**, Materials and Methods).

RF models were trained to predict PPIs in *E. coli* K12 using MIWSCs computed from the SVD spectrum of ***D^domain^*** (MIWSCsdomain). These models were validated in the same multilevel classification tasks as described above for ***D^OGG^*** (**Figure 3 - figure supplement 2**). When compared to RF models trained on features of existing computational and experimental methods, the RF models trained on MIWSCsdomain ranking 1st, 1st, and 3^rd^ for the classes of not-interacting, indirect PPI, and direct PPI respectively (**Table S6**).

Taken together, these results illustrate that spectral decomposition of orthologous gene content across bacterial proteomes separates covariation arising from different biological scales regardless of whether orthology is defined through orthologous gene groups or protein domains. As a result of this spectral separation, spectral correlations computed across sets of SVD components of OGG or domain covariation can produce accurate predictions of PPIs at different biological scale, i.e. indirect PPIs and direct PPIs.

### A statistically-defined hierarchy of protein interaction networks describing the emergent ***phenotype of directed motility in E. coli K12***

Understanding the molecular basis of a phenotype requires (i) identifying units of collective function at different biological scales and (ii) relating these scales to create a hierarchical model of how a phenotype emerges from a set of proteins. A useful example is the experimentally derived hierarchical model of directed motility in *E. coli* K12 (KEGG hierarchy, BRITE ECO:02035). At the lowest levels in this hierarchy, physical interactions between proteins create small units of collective structure and function, such as a basal body, rod, ring, motor, and filament. Integration of these structures and their individual functions produces the flagellum, a machine that turns to move the cell. Integration of the flagellum and the chemotaxis system ultimately produces directed motility – the ability to move purposefully along a chemical gradient.

We hypothesized that we could derive a multiscale, hierarchical model of this phenotype in a purely data-driven and unbiased manner using only the SVD spectrum of ***D^OGG^***. To do so, we first developed a model of spectral correlations between non-interacting proteins. We then applied this model to identify ‘significant’ protein correlations within different regions of the SVD spectrum. These significant proteins correlations represented statistically predicted protein interactions. Next, we defined a metric termed ‘spectral depth’ as the deepest spectral position to which two proteins remained significantly correlated. We posited that applying serial thresholds to spectral depth would identify a tree-like hierarchy where the root of the tree is the protein interaction network observed at shallower spectral depth thresholds and the branches are the networks defined at deeper spectral depth thresholds. The details of creating the model of non-interacting protein correlations and defining a hierarchy using spectral depth are detailed below.

To define a model of spectral correlations between non-interacting proteins, we first considered the distribution of all pairwise spectral correlations centered on SVD1000 for the proteins encoded in the proteome of *E. coli* K12. Our rationale was that since the vast majority of proteins do not interact to produce a collective function, the distribution of all-by-all spectral correlations approximates that of correlations between proteins that do not functionally interact. We observed that the variance of this distribution decreased rapidly as the correlation window widened until about a 100 component width – motivating our choice of computing correlations over sets of 100 components (**Figure 4 – figure supplement 1A,B**). We computed distributions of all-by-all correlations between *E. coli* K12 proteins across windows centered on different regions of the SVD spectrum and found them to be superimposable (**Figure 4 – figure supplement 1C**). Additionally, we computed such distributions for proteins from other diverse bacteria and found them to be superimposable with those derived from *E. coli* K12 (**Figure 4 – figure supplement 1D**). These properties enabled us to define a constant threshold for significant spectral correlations between two proteins across any 100 component SVD window. The p-value derived from the empirical CDF of this model decreased rapidly until a threshold value of 0.29 (**Figure 4 – figure supplement 1E**). Therefore, we chose the value of 0.29 associated with a p-value of 0.018 as the threshold of spectral correlations signifying functional interactions between proteins derived from any bacterial proteome within any region of the SVD spectrum.

**Figure 4.**
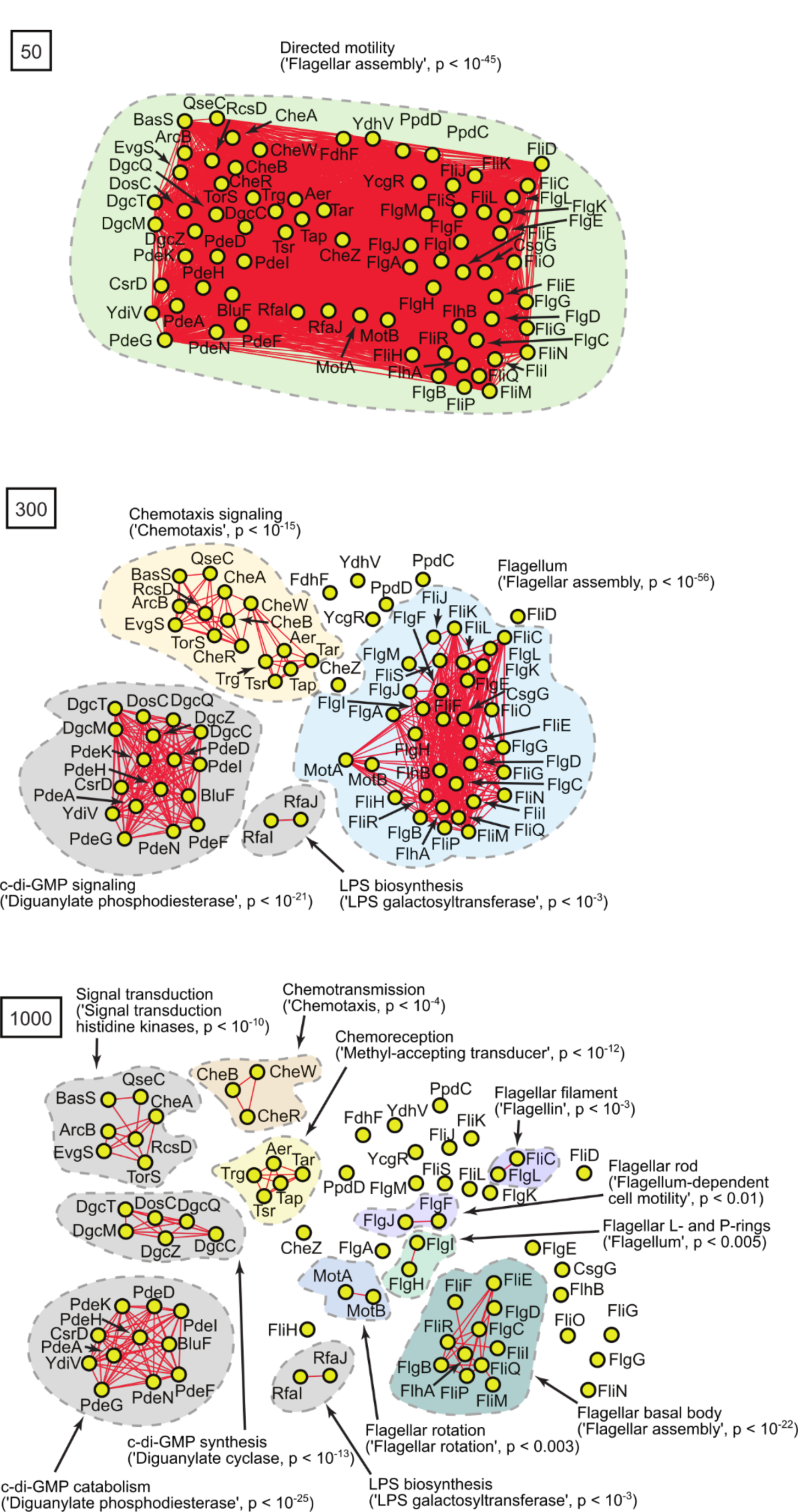
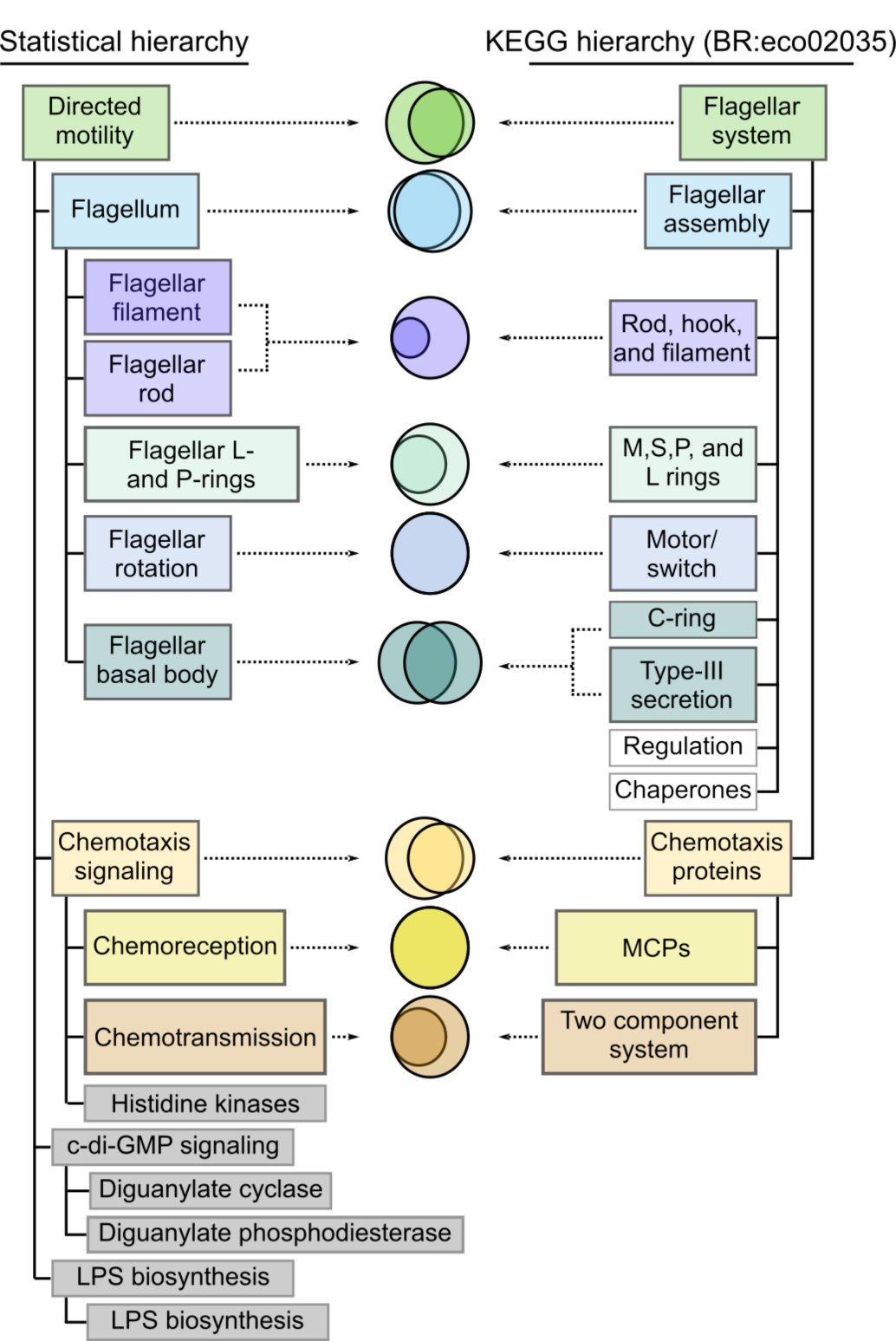
A hierarchical model of *E. coli* K12 motility derived by serially thresholding ‘spectral depth’ resembles that described in the KEGG database. (A) The model contained 75 proteins that were identified as significantly correlated with FliC across SVD34 to SVD134.Statistical interaction networks were defined by thresholding spectral depth at 50 (top panel), 300 (middle panel), and 1000 (bottom panel). Nodes (yellow circles) represent proteins and edges (red lines) represent statistical interactions. Shaded contours identify discrete subnetworks and are labeled with their assigned function based on interpretation of gene-set enrichment analysis (GSEA) and literature review. The most significantly enriched ontological term produced by GSEA and the associated p-value is shown in parentheses for each subnetwork (**Table S7**). (B) Comparison of the statistically-derived model (left) to the KEGG model (BR:eco02035, right) of *E. coli* K12 motility. Venn diagrams represent the overlap between the sets of proteins in the indicated subnetwork of panel A (left) and the indicated KEGG category (right).

To develop a hierarchical model of the directed motility phenotype in *E. coli* K12 we first identified all proteins (n = 75) that were significantly correlated with FliC, the flagellar filament protein, over the first spectral window enriched for non-phylogenetic information (SVD34-SVD134) (**Figure 4 – figure supplement 2A**). For these proteins, we computed pairwise spectral correlations across all 100 component windows of the SVD spectrum of ***D^OGG^***. We observed that some pairs (e.g. MotA and MotB) remained significantly correlated as the spectral window was shifted deep into the SVD spectrum while other pairs (e.g. MotA and CheR) were significantly correlated only across the shallower regions of the SVD spectrum (**Figure 4 – figure supplement 2B**). We computed the position at which the pairwise correlation first dropped below the significance threshold defined by our model of correlations between non-interacting proteins. We define this position as the ‘spectral depth’ of correlation. We computed the spectral depth for all pairs of proteins that were significantly correlated with FliC across SVD34 to SVD134 (**Figure 4 – figure supplement 2C**). Apply a threshold to spectral depth generates an adjacency matrix where a pixel value of ‘1’ indicates a pair of proteins that share a spectral depth that is as deep or deeper than the threshold value (**Figure 4 – figure supplement 2D**).

This adjacency matrix can be used to construct a protein interaction network at the thresholded spectral depth.

We constructed protein interaction networks from the adjacency matrices produced by applying spectral depth thresholds of 50, 300, and 1000 (**Table S7**). At a spectral depth of 50, we observed a single densely connected network devoid of obvious substructure (**Figure 4A**, top panel). Gene set enrichment analysis (GSEA) indicated that this network was enriched for functional terms related to ‘flagellar system’ (p < 10^-45^) (Huang *et al*., 2009; Huang *et al*., 2009, Materials and Methods). Progressing to spectral depth of 300, we observed that the network at 50 fractured into four discrete subnetworks (**Figure 4A**, middle panel). These subnetworks were significantly enriched for terms related to ‘Chemotaxis signaling’ (p < 10^-15^), ‘Flagellum’ (p < 10^- 56^), ‘LPS biosynthesis’ (p < 10^-3^), or ‘cyclic di-GMP signaling’ (p < 10^-21^). Progressing to spectral depth of 1000, the subnetworks at 300 fractured further yielding 9 discrete subnetworks. Each subnetwork was significantly enriched for terms related to a specific function such as ‘cyclic di-GMP catabolism’ (p < 10^-25^) and ‘cyclic di-GMP synthesis’ (p < 10^-13^) or ‘chemotransmission’ (p < 10^-4^) and ‘chemoreception’ (p < 10^-12^) (**Figure 4A**, bottom panel).

Taken together the three network diagrams derived at spectral depths 50, 300, and 1000 depict a hierarchy of structure and function. Subnetworks observed at deeper spectral depths integrate to form the subnetworks observed at shallower spectral depths. As the subnetworks coalesced, the p-value associated with GSEA remained highly significant while the ontology of the significantly enriched terms changed. We interpret these observations to mean that as we ascend the statistical hierarchy, molecular descriptions of new biological functions emerge from the integration of functional units at lower levels.

We compared our hierarchical model with the model of *E. coli* K12 motility detailed within the KEGG database (BR:eco02035) (Kanehisa *et al*., 2017) (**Figure 4B**). The two models were similar in several ways. First, 44 of 55 of the proteins listed in the KEGG hierarchy also appeared in the statistical hierarchy. Second, 7 of the 12 categories listed in the KEGG hierarchy had a one-to-one correspondence with a subnetwork of the statistical model sharing an overlapping set of proteins and similar descriptive label. Finally, both hierarchies shared a conserved architecture consisting of the integration of chemoreception and chemotransmission into chemotaxis signaling, the integration of flagellar substructures into the flagellum, and at the most global level the integration of chemotaxis and the flagellum. The most striking difference was that our statistical hierarchy included subnetworks related to cyclic-di-GMP signaling and LPS biosynthesis which were absent from the KEGG hierarchy. Prior experimental studies have provided direct genetic evidence that these systems are involved in *E. coli* K12 motility (Paul *et al*., 2010, Walker *et al*., 2004).

Overall, of the 75 proteins in our hierarchical model of *E. coli* K12 motility, 44 (59%) were represented in the KEGG hierarchy, 28 (37%) were missing from the KEGG hierarchy but supported by prior experimental evidence in the literature, and only 3 (4%) remained unvalidated (CsgG, PpdD, TorS) (**Table S7**). Taken together, these results illustrate that identifying the *E. coli* K12 proteins that were significantly correlated with FliC and then serially thesholding their spectral depth produced a valid multiscale, hierarchical model of *E. coli* K12’s directed motility phenotype.

### Robustness and generalizability of defining statistical hierarchies using spectral depth

We performed four additional to test the robustness and generalizability of our approach. First, we characterized motility in *E. coli* K12 using MotB, the flagellar motor protein, as a query. We found a similar hierarchical architecture as observed using FliC as the query with chemotaxis signaling, flagellum, and cyclic-di-GMP signaling modules appearing at spectral depth 300, and more fine-grained subnetworks appearing in deeper layers (**Figure 4 – figure supplement 3**, **Table S8**). To test generalizability across organisms, we created a model of motility in *Bacillus subtilis* 168 using its flagellar filament protein as a query (Hag). This analysis again produced a hierarchical model of motility that (i) recapitulated the corresponding KEGG hierarchy, (ii) identified proteins missing from the KEGG hierarchy that are known effectors of *B. subtilis* motility, and (iii) identified a small number of putative motility effectors (**Figure 4 – figure supplement 4**, **Table S9**). Next, we tested if our method could generalize to non-physically coupled pathways. We produced a model of amino-acid metabolism in *E. coli* K12 using the query protein HisG, an enzyme involved in Histidine biosynthesis. The resultant hierarchical model identified 130 proteins that were densely connected at spectral depth 50. Progressing to deeper spectral depths revealed modules corresponding to specific functions, such as amino acid and nucleotide biosynthesis. At yet deeper spectral depths, modules enriched for proteins involved in the synthesis of specific amino acids became evident (**Figure 4 – figure supplement 5**, **Table S10**). Finally, we demonstrated that valid statistical models of *B. subtilis* 168 and *E. coli* K12 motility could be derived by serially thresholding spectral depth of correlations within the SVD spectrum of ***D^domain^*** (**Figure 4- figure supplement 6**, **Table S11**, **Table S12**). Taken all together, these analyses demonstrated that serially thresholding spectral depth produces a hierarchical model of biological pathways across different query proteins, organisms, pathways, and orthologous features.

### Using the structure of a statistically defined hierarchy to aide in the discovery of novel ***genotype-phenotype relationships***

The hierarchical models produced by serially thresholding spectral depth recapitulated the known architecture of several well-studied biological phenotypes without incorporating any prior knowledge of biological organization. This motivated the hypothesis that these models could also reveal new biological organization that was not previously appreciated. We tested this idea by generating a hierarchical model of motility in *Pseudomonas aeruginosa*, using it to assign both a general and specific function to a previously uncharacterized protein, and experimentally validating these predictions.

*P. aeruginosa* is uses two different types of motility – propulsive motility based on a flagellum and twitch motility based on a pilus (Kearns *et. al.*, 2001; Rashid and Kornberg, 2001). Using PilA, a structural component of the pilus, as a query we identified a network of 141 proteins as significantly correlated across SVD34 to SVD134. We produced network configurations for these proteins using spectral depth thresholds of 50 and 300 (**Table S13**). At a spectral depth of 50 a single densely connected network was observed (**Figure 5 – figure supplement 1A**). Significantly enriched terms for this network were ‘methyl-accepting chemotaxis’ (p < 10^-34^), ‘cell motility and secretion’ (p < 10^-33^), ‘two-component system’ (p < 10^- 27^), ‘type IV pilus-dependent motility’ (p < 10^-10^), and ‘flagellar assembly’ (p < 10^-6^), suggesting that these proteins are collectively involved in the global function of directed motility. At spectral depth 300, the network fractured into 18 different discrete subnetworks that were enriched for specific functions (**Figure 5A**, **Figure 5 – figure supplement 1B**). The largest subnetworks were enriched for ‘methyl-accepting chemotaxis protein’ (p < 10^-59^), ‘pilus motility’ (p < 10^-17^), or ‘bacterial flagellum’ (p < 10^-21^). We noted four proteins in the pilus subnetwork (Q9I5G6, Q9I5R2, Q9I0G2, Q9I0G1) that were annotated as ‘uncharacterized protein’ in UniProt (**Figure 5B**). Further review of STRING, GO, BIOCYC, and PFAM revealed only that Q9I5G6 contains a ‘domain of unknown function’ (DUF4845). Based upon their membership in the pilus subnetwork at spectral depth 300, we hypothesized that these proteins may contribute to the general function of directed motility by affecting the specific function of pilus-based motility. Furthermore, the lack of connections to the flagellum subnetwork suggested that these proteins would not impact flagellar based motility.

**Figure 5.**
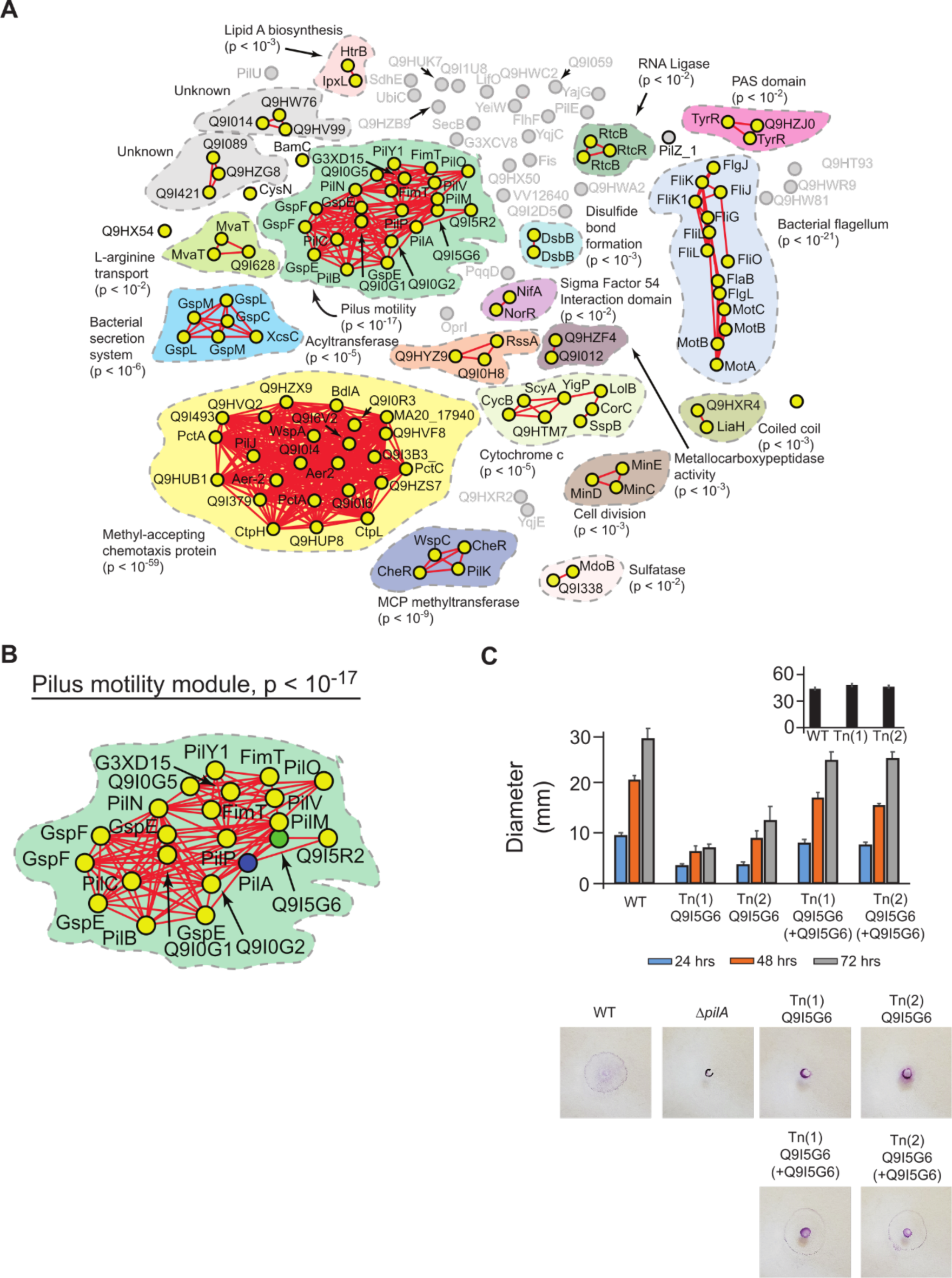
Prediction and experimental validation of a novel effector of twitch-based motility in P. aeruginosa (PAO1). (**A**) Statistical network derived by applying a spectral depth threshold of 300 to the set of 141 protein in *P. aeruginosa* that were significantly correlated with PilA across SVD34 to SVD134. Nodes (yellow circles) represent proteins and edges (red lines) represent statistical interactions. Shaded contours identify statistical subnetworks and are labeled with their assigned function based on interpretation of gene-set enrichment analysis (GSEA) and literature review. The most significantly enriched ontological term produced by GSEA and the associated p-value is shown in parenthesis for each subnetwork (**Table S13**). (**B**) The pilus motility subnetwork from panel A. Nodes representing PilA and Q9I5G6 are colored blue and green respectively. (**C**) Time-course of pilus-based motility for parent (WT), two transposon mutants of Q9I5G6 (Tn(1) Q9I5G6, Tn(2) Q9I5G6), and transposon mutants complemented with Q9I5G6 (Tn(1) Q9I5G6 +Q9I5G6, Tn(2) Q9I5G6 +Q9I5G6). Inset shows results of flagellar motility for the parent strain (WT), and the two transposon mutants of Q9I5G6 (Tn(1), Tn(2)) 24 hours post-inoculation. Representative images of the crystal-violet stained plates are shown.

To test these predictions, we screened single-gene transposon mutants of *P. aeruginosa* (PAO1) for twitch-based or flagellar-based motility using established experimental assays (Materials and Methods) (Kearns *et. al.*, 2001; Rashid and Kornberg, 2001, Little *et. al.*, 2018). Transposon mutants of Q9I5R2, Q9I0G2, and Q9I0G1 exhibited motility that was not significantly different from the parent strain in both assays (**Table S14**). In contrast, we found that two different transposon mutants of Q9I5G6 exhibited significantly reduced twitch motility velocity over 24, 48, and 72 hours compared to the parent strain (**Figure 5C,** p < 10^-4^ by Dunnett’s multiple comparisons test). This phenotype resembled that of a knockout strain of pilA and was reversed upon trans-complementation. In contrast, flagellar-based motility of Q9I5G6 was not significantly different from that of the parent strain (**Figure 5C** – inset, p > 0.05). These results illustrate that Q9I5G6 is a previously unappreciated effector of the global directed motility phenotype in *P. aeruginosa* that specifically impacts twitch-based motility. As such, these experiments provide a proof of concept of how our hierarchical models may aid in discovering novel genotype-phenotype relationships.

## Discussion

Connecting genotype to phenotype is a central goal in biology. Achieving this goal requires understanding how the collection of proteins in a proteome interact at different scales spanning protein complexes, pathways, and cellular phenotypes. Here, we have shown that a hierarchy of protein interaction networks can be extracted from analyzing covariation across an ensemble of bacterial proteomes. Key to this outcome were three important results. First, when we spectrally decomposed proteome variation using SVD we found that biological information mapped onto the SVD spectrum in a specific way: shallow components were enriched for phylogenetic relationships, deeper components for functional interactions between proteins in pathways, and even deeper components for physical interactions within protein complexes.

Second, we found that spectral correlations measured across sets of SVD components defined features that informed accurate classification of protein pairs as non-interacting, indirect PPI, or direct PPI. Third, we developed the concept of computing a spectral depth of correlation and found that serially thresholding spectral depth produced a hierarchical model of protein interaction networks. These models closely resembled the known hierarchical organization for several well-studied bacterial phenotypes. Finally, we illustrated the utility of generating these unbiased hierarchical models by developing a model of motility in *P. aeruginosa* and using it to predict global and local functions for a previously uncharacterized protein.

We call our approach SCALES—Spectral Correlation Analysis of Layered Evolutionary Signals. To facilitate access to SCALES and other methods described in this paper, we have developed (i) a precomputed database of proteome-wide indirect (122,725,727) and direct (19,546,063) PPI predictions for all 7,047 UniProt reference bacterial proteomes; (ii) a tool for predicting indirect and direct PPIs for a user-input proteome; (iii) a tool for generating and interrogating a hierarchical model for a query protein of interest using SCALES. All of these can be found at scales.cri.uchicago.edu

### Discarding global components of covariation purifies PPI information

Variation in bacterial proteomes arises from different sources of information and noise. An established approach to separating information from noise is to spectrally decompose the variation using SVD and then to identify which SVD components can explain more of the total variance than a random process (Wigner, 1967). This leads to considering the *k* most global SVD components (Franscescini *et. al.*, 2016). In contrast, in our study, we empirically mapped the distribution of biological information across the entire SVD spectrum. Our results did not match our initial expectation that the most global components would inform the highest fidelity PPI predictions and minor components would solely contain noise. Instead, we found that global components of covariation primarily reflected phylogeny, and PPI predictions based on these components were low quality. On the other hand, we found that excluding global components while retaining minor components harboring a minuscule amount of variance produced high-fidelity PPI predictions. We interpret these results to mean that percent variance per component does not indicate ‘importance’ of biological signal and discarding major components of covariation may actually purify functionally relevant information.

### Spectral depth: a metric that extracts a hierarchy from the SVD spectrum

SVD sequentially defines orthonormal vectors (components) that maximize the compression of the remaining unexplained data variance. We found that there is a direct mapping between the position of a component within the SVD spectrum and level in the hierarchy of biological organization (**Figure 1C**). Likely this mapping reflects intrinsic differences in the compressibility of biological variation arising from different hierarchical levels spanning protein complexes, pathways, and phenotypes. However, SVD alone does not reveal how the different levels in the hierarchy are related. Therefore, to extract a model of how protein interactions are hierarchically organized to generate a phenotype, we devised the metric of spectral depth, the tracking of the persistence of correlations across the SVD spectrum. We found that this metric enables predicting the integration of PPIs into complex structures approximating pathways and phenotypes.

### Limitations

There are several limitations to the methods developed in this manuscript related to: 1) the feature selection, 2) data requirements, and 3) lack of mechanistic insights. We discuss these limitations in the following paragraphs.

We observed two major limitations related to the definition of an orthologous feature.

First, there are proteins that have no annotated conserved protein domain or OGG. For example, as many as 12.3% and 16.3% of the *E. coli* K12 proteins lacked a domain or OGG annotation respectively. These proteins cannot be assigned to interactions or units of function by our methods. Second, there are proteins that share an infinite spectral depth of significant correlation because they contain redundant annotations. These proteins may or may not contribute to a shared biological function. We anticipate that continued expansion of available bacterial genome sequences will improve ortholog annotation and help alleviate these limitations. In addition, phylogenomics may help by providing methods that can capture orthology relationships spanning both short and long timescales of evolution (Nagy *et. al.*, 2020. Despite the current limitations, the sets of interactions we predicted were more accurate and complete when benchmarked against existing methodologies (**Figure 2**).

Recently Cong and colleagues reported a method for inferring PPIs using amino-acid level coevolution (Cong *et al*., 2019). Although we did not incorporate high-resolution amino-acid information into our method, we observed that our method produced more precise and complete PPI predictions in *M. tuberculosis* (**Figure 2D**). At this time, it is unclear if these performance differences were related to the feature of observation or methodological differences. However, the results of our comparative analysis suggest that merely increasing the resolution of genomic feature does not necessarily equate to more accurate PPI prediction.

More work will be required to determine if incorporation of amino-acid variation could help resolve spectral correlations between proteins with overlapping low-resolution orthologous features. Nonetheless, one important implication of using a lower resolution feature, like OGGs, is our ability to compute proteome-wide PPI predictions in a matter of minutes.

To what degree is the ability to recover a hierarchy of biological organization dependent on the ensemble of proteomes? We reason that there are two important characteristics of the ensemble—the number and diversity of the proteomes. With respect to the first, we observed that thousands of SVD components were required to provide a protein complex level description of the 7,047 UniProt bacterial reference proteomes likely reflecting the poor compressibility of this granular information. SVD can only define as many components as the smallest dimension of the matrix under interrogation. In the era of genomic sequencing the number of biological replicates (the rows) is typically more limiting than the number of features (the columns). As such, our ability to recover protein complex level information depended on having thousands of available proteomes. With respect to the second, the ensemble we used was non-redundant, representing a diversity of species (**Figure 1 – figure supplement 1**). Why was this facet important? Consider the extreme case where all rows represent proteome sequences from the same organism. In this case, all statistical variance would be contained within a single SVD component, precluding the ability to separate and relate different levels of hierarchical organization. In the future, it may be useful to explore using the shape of the SVD spectrum to guide sequencing efforts so that the number as well as diversity of replicates within the ensemble are sufficient.

A notable characteristic of the methods developed here is that they are inherently ‘mechanism-free’, a quality we view as both a limitation and a strength. It is a limitation in that our approach can identify interactions but cannot provide insight into the function or nature of those interactions. However, it is a strength because we are not limited by prior experimental results and methods. As such, we believe that our approach may powerfully guide discovery of novel biology.

### The potential of generalizing SCALES to other biological systems

As ‘big-data’ in biology is becoming increasingly commonplace, to what degree are the approaches developed here applicable to other biological systems? We note that the spectral properties of any given dataset will be unique. As such, re-application of these methods to a new dataset will require following the steps outlined in this work: creating an ensemble, identifying relevant benchmarks, mapping the benchmarks onto SVD components, and developing a model of random spectral correlations to define spectral depth. Beyond these practical considerations, a larger question is whether the methods described here are appropriate for interrogating biological systems more generally. SCALES represents a statistical way to describe emergence—the integration of individual components into layers of collective units of function. The property of emergence spans many biological systems, from proteins to ecosystems. Thus, while it remains to be tested, it may be true that SCALES is a generally useful approach to learning the hierarchical architecture of biological systems.

## Materials and Methods

### Generating **D^OGG^**

All bacterial proteomes (n = 7,047) in the 2020_02 release of the Uniprot Reference Proteome database were downloaded on 05/20/2020 (**Figure 1 – figure supplement 1**) (The UniProt Consortium, 2019). OGGs were annotated using eggNOG-mapper V2 at the level of bacteria (‘@2’) (Huerta-Cepas *et al*., 2017; Huerta-Cepas *et al*., 2019). An OGG count matrix was assembled (***D^OGG^***, **Figure 1A**) where rows were defined as proteomes, columns were defined as OGGs, and the value in each cell was the number of annotations an OGG in a proteome. The number of annotations was used to preserve as much information as possible versus the strategy of considering binary occurrence. All OGGs present in fewer than 1% of the proteomes were removed leaving 10,177 unique columns in ***D^OGG^***.

### Singular value decomposition (SVD) of D^OGG^

Singular Value Decomposition (SVD) was performed on ***D^OGG^***. First, the raw data matrix was centered and standardized by

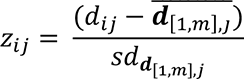

where *z*_*ij*_ is the *ij*^th^ element of the z-scored matrix ***Z^OGG^***, *d*_*ij*_ the *ij*^th^ element in the initial data matrix 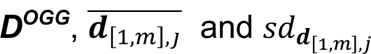 are the mean and standard deviation of the *j*^th^ column vector of ***D^OGG^***, and *m* is the total number of rows in ***D^OGG^*** (*m* = 7,047). ***Z^OGG^*** was factorized using SVD:

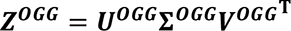

***U^OGG^*** is an m x K matrix where rows are proteomes, columns are the ‘left singular vectors’, and each element is the projection of a proteome onto a left singular vector. ∑^OGG^ is a m x n diagonal matrix where the K non-zero diagonal entries are ‘singular values’ and decrease in magnitude with position along the diagonal. ***V^OGG^*** is a n x K matrix where the rows are OGGs, columns are the ‘right singular vectors’, and each element is the projection of an OGG onto a right singular vector. m and n are the number of rows (m = 7,047) and columns (n = 10,117) in ***Z^OGG^***. K is the total number of SVD components (K = 7.047). The fraction of data variance explained by SVD component k is computed through the following equation:

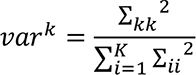

where var^k^ is the fractional variance explained by the *k*^th^ SVD component, Σ_*kk*_ and Σ_*ij*_ are the *k*^th^ and *i*^th^ singular values respectively, and *K* is the total number of singular values. The plot of fractional variance per component versus component number are shown for ***D^OGG^*** in **Figure 1 – figure supplement 2D**.

### Assembling benchmarks that collectively represent the hierarchy of biological organization

The various benchmarks described within this section can be found in **Table S2, S3**.

### Phylogeny benchmarks

NCBI phylogenetic strings were mapped to the NCBI taxonometric IDs for each of the 7,047 bacteria represented in ***D^OGG^*** using taxonkit 5.0 (https://bioinf.shenwei.me/taxonkit/) on 5/20/2020. Five different benchmarks were generated corresponding to pairs of proteomes that share identical phylogenetic substrings down to the level of phylum (n = 5,841,696), class (n = 2,460,194), order (n = 807,338), family (n = 434,753), or genus (n = 267,794).

### STRING Nonbinding benchmark

STRING database annotations were downloaded for the *Escherichia coli* K12 proteome (STRING ID 511145) on 7/22/2019. A benchmark was assembled to include all protein pairs (n = 14,793) with a nonzero combined STRING score that did not share a ‘binding’ action annotation. This benchmark was expected to be enriched for indirect protein-protein interactions (PPIs).

### GO terms benchmark

Biological function’ GO term annotations were mapped for the 4,391 proteins in the *E. coli* K12 proteome through the UniProtKB API. A benchmark was assembled containing the 79,794 protein pairs that share at least 1 GO term annotation. This benchmark likely contained a mixture of indirect and direct PPIs.

### STRING benchmark

STRING database annotations were downloaded for the *E. coli* K12 proteome (STRING ID 511145) on 7/22/2019. A benchmark was assembled comprised of all (n = 20,216) protein pairs with a nonzero combined STRING score. This benchmark included a mixture of pairs with (n = 14,793) and without (n = 5,423) a ‘binding’ annotation and therefore is presumed to contain a mixture of direct and indirect PPIs.

### ECOCYC benchmark

A previously published benchmark included 915 pairs of *E. coli* K12 proteins selected from the set of complexes in the ECOCYC database after intentionally excluding large complexes with greater than ten proteins to enrich for directly interacting pairs of proteins (Cong *et al*., 2019, Keseler *et al*., 2017). This benchmark is assumed to primarily represent direct PPIs.

### Coev+ benchmark

A previously published set of 1,600 direct PPIs in *E. coli* K12 identified by a hybrid method combining the results of amino acid coevolution (AA Coev) and prior experimental data (Cong *et al*., 2019).

### PDB benchmark

A previously published set of 809 direct PPIs in *E. coli* K12 selected by the criteria that they, or closely homologous proteins, have been observed to interact in a crystal structure in the PDB (Cong *et al*., 2019).

The type of biological information reflected in each benchmark is summarized in **Figure 1B**.

### Computing protein-protein spectral correlations

A row vector in the matrix ***V^OGG^*** contains the projections of a single OGG onto each of the SVD components as described by:

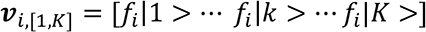

where *v*_i, [*i,k*]_ is the *i*^th^ row vector of the matrix ***V^OGG^***, *fi* is the OGG represented in row *i* of matrix ***V^OGG^***, *f*_*i*|*k* > is the projection of *fi* onto the SVD component *k* (1 ≤ *k*_ ≤ *K*), and *K* is the total number of SVD components.

A vector representing the projections of a protein onto the SVD components was generated by averaging the vectors corresponding to the projections of all OGGs annotated within the protein:

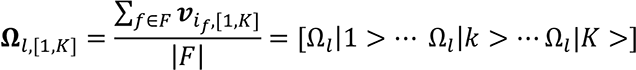

where Ω_*l*,[1,*K*]_ is the vector of averaged projections for all OGGs in protein *l* onto the *K* SVD components and is used as a surrogate for the projections of protein *l* onto the *K* SVD components, *f* is an OGG in *F* (the set of OGGs encoded in protein *l*), *if* is the index of the row in the matrix ***V^OGG^*** that contains *v*_i,f,[1, *k*]_ (the vector of projections of OGG *f* onto the *K* SVD components), |*F|* is the number of OGGs in *F*, *k* is a single SVD component (1 ≤ *k* ≤ *K*), Ω_1_ |*k* > is the average projection of the OGGs in protein *l* onto component *k*, and *K* is the total number of SVD components. An example of this process is illustrated in **Figure 1 – figure supplement 4A-F**.

A subvector of Ω_*m*,[*a,b*]_ was extracted so as to only consider the projections of protein *l* onto a window of SVD components as described by:

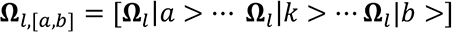

where Ω_*l*,[*a,b*]_ is a vector of the projections of protein *l* onto the set of SVD components ranging from component *a* to component *b*, 1 ≤ *a* ≤ *K*-1, and 2 ≤ *b* ≤ *K*.

The correlations between protein *l* and protein *m* within a spectral window was defined

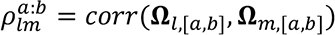

where 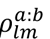 is the correlation between proteins *l* and *m* within the spectral window ranging from SVD component *a* to SVD component *b, corr* denotes the Pearson correlation, and Ω_*l*,[*a,b*]_ and Ω_[*a,b*]_ are the vectors containing the projections of proteins *l* and *m* onto the SVD components within the spectral window respectively. An example of this process is illustrated in **Figure 1 –**figure supplement 4G.

A model of random spectral correlations was generated by row shuffling the matrix ***V^OGG^*** and then computing protein-protein spectral correlations as above (**Figure 1 – figure supplement 5**).

### Computing Mutual Information (MI) between spectral correlations and benchmarks of biological knowledge

For each phylogenetic benchmark, one-hundred bootstraps were generated consisting of equal numbers of randomly selected pairs of proteomes that do or do not share an identical phylogenetic substring annotation in the benchmark. For each protein interaction benchmark, one-thousand bootstraps were generated consisting of equal numbers of randomly selected pairs of proteins that do or do not share an interaction annotation in the benchmark. For bootstraps of both phylogenetic and protein interaction benchmarks, the number of pairs sharing an annotation was equal to the number of pairs indicated for each respective benchmark in the section

### ‘Assembling benchmarks that collectively represent the hierarchy of biological organization’

Spectral correlations across all 5-component windows of the SVD spectrum between component 1 and component 3000 were computed for the proteome or protein pairs in each bootstrap. The uncertainty surrounding the spectral correlations within a single window was calculated as the differential entropy (Cover and Thomas, 2006):

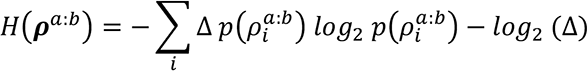

where *ρ*^*a:b*^ is the vector of spectral correlations over the window ranging from components *a* to *b* for the pairs in the bootstrap, H*ρ*^*a:b*^J is the differential entropy of *ρ*^*a:b*^ , *ρ*^*a:b*^ is a bin of correlation values produced by quantizing the continuous-valued correlations in *ρ*^*a:b*^ , *p*(*p*_*i*_^*a:b*^) is the probability of observing a correlation value within *ρ*^*a:b*^, and Δ is the width of the quantization bins. In the present study Δ = 0.25.

Uncertainty surrounding spectral correlations given knowledge of the phylogenetic relationships or protein interactions annotated within a benchmark is described by the conditional entropy:

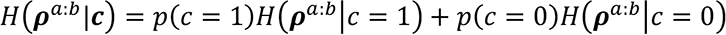

where *c* is a binary vector that assumes a value of 1 or 0 if a pair of proteomes or proteins do or do not share an annotation in the corresponding benchmark respectively, H*ρ*^*a:b*^ |*c* is the uncertainty surrounding the windowed spectral correlations given knowledge of *c*, *p*(c=1) and *p*(c=0) are the probability of a l or 0 in *c* respectively, and H*ρ*^*a:b*^ | *c* = 1) and H*ρ*^*a:b*^ | *c* = 0) are the uncertainties surrounding the spectral correlations for subsets of pairs in the bootstrap that correspond to a value of 1 or 0 in *c* respectively and are computed using the differential entropy equation described above.

Finally, MI was computed as the difference between the uncertainty surrounding the spectral correlations with and without knowledge of the benchmark:

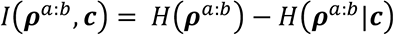

where *Iρ*^*a:b*^ , *c*J is the estimate for the MI shared between the spectral correlations and the benchmark. A model of random MI was generated by computing the MI shared between the spectral correlations within row-shuffled versions of ***U^OGG^*** or ***V^OGG^*** and the benchmarks of phylogeny and protein interactions respectively (**Figure 1 – figure supplement 5**). The distributions of MI estimates for the different benchmarks arising from the data or random model are summarized in **Figure 1 – figure supplement 6**.

### Calculation of MI cumulative distribution functions (cdfs) shown in **Figure 1C**

Each point in the MI cdfs shown in **Figure 1C** was computed as (for the window centered on component *w* of the SVD)

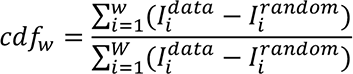

where *cdfw* is the value of the cdf at spectral position *w, Ii^data^* is the MI observed in window *i*, *Ii^random^* is the MI produced by random correlations in window *i*, *W* is the total number of windows, and 1 ≤ *w* ≤ *W*. Because we considered 5-component spectral windows within the first 3000 components, *W =* 2997.

### Training and validating Random Forests (RF) models for predicting PPIs in E. coli K12 using MIWSCs

#### Assembling a ‘gold-standard’ dataset

A ‘gold-standard’ dataset for *E. coli* K12 PPIs was assembled and consisted of 72,000 not- interacting, 1,226 indirect PPIs, and 72 direct PPIs. All pairs defined as ‘direct PPI’ satisfied three criteria: they shared amino-acid level coevolution (Coev+ benchmark), were annotated in the same protein complex in the ECOCYC benchmark, and interacted in the PDB benchmark. All indirect PPIs were selected based on the following criteria: they shared a ‘non-binding’ type interaction annotation in the STRING Nonbinding benchmark, shared a ‘biological function’ interaction in the GO benchmark, and did not share an interaction annotation in any of the benchmarks of direct PPIs (Coev+, ECOCYC, or PDB). The ‘not-interacting’ pairs did not share an interaction annotation in any of the benchmarks (GO, STRING Nonbinding, STRING, Coev+, ECOCYC, or PDB). The not-interacting set was subsampled to exceed the number of physically interacting pairs by 1000-fold (Rajagopala *et al*., 2014; Cong *et al*., 2019).

The gold standard pairs were randomly partitioned into training (60%) and validation (40%) datasets. Fifty such random partitions were generated to assess the reproducibility of the results of the machine-learning task described below.

#### Training RF models

RF models consisting of 100 decision trees were trained to classify pairs of proteins in *E. coli* K12 as not-interacting, indirect PPIs, or direct PPIs by feeding the labeled training set examples to the TreeBagger algorithm (Matlab, v2020a). This process was repeated for each random partition of the gold-standard dataset yielding an ensemble of 50 RF models per feature.

#### Validating RF models using the validation dataset

Each trained RF model was subjected to three validation tasks of classifying interaction types for unlabeled pairs of *E. coli* K12 proteins in the validation portion of the gold-standard (40%) (**Figure 2 – figure supplement 1, Figure 2 – figure supplement 2A**). The model performance was evaluated by computing an F-score for each interaction type (not-interacting, indirect PPIs, direct PPIs), where F-score is the harmonic mean of precision and recall, precision is the ratio of the number of correctly predicted interactions within a class to the total number of predicted interactions in a class, and recall is the ratio of the number of correctly predicted interactions within a class to the total number of interactions of that class.

### Training and validating RF models on quantitative features of existing methods

For each feature extracted from existing methods described below, RF models were trained and validated using the identical protocol as for MIWSCs (described in the section *<u>Training and validating Random Forests (RF) models for predicting PPIs in E. coli K12 using MIWSCs</u>*).

#### Existing experimental features

Previously published datasets derived from large scale experimental PPI screens in *E. coli* K12 were used to generate a set of four different experimental features including: gene interaction scores from a gene epistasis screen (Epistasis, n = 41,820), sum log-likelihood scores from an affinity purification mass spectrometry screen (APMS1, n = 12,801), protein interaction scores from an affinity purification mass spectrometry screen (APMS2, n = 291), and binary pairs from a yeast-two hybrid screen (Y2H, n=1,766) (Rajagopala *et al*., 2014; Babu *et al*., 2014; Babu *et al*., 2018; Hu *et al*., 2009).

#### Existing computational features

Gene co-occurrence, gene fusion, and gene neighborhood subscores for *E. coli* K12 (STRING ID 511145) were extracted from the STRING database on 7/22/2019 (Szklarczyk *et al*., 2019; Rajagopala *et al*., 2014; Babu *et al*., 2014; Babu *et al*., 2018; Hu *et al*., 2009). Any pairs without an interaction annotation in the STRING database were assigned a subscore of zero.

#### Binary MI (b-MI) feature

The b-MI feature was modeled after the popular phylogenetic profiling method of Pelligrini and colleagues (Pellegrini *et al*., 1999). First, a binary OGG content matrix was definedas follows:

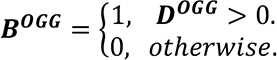

Where ***B^OGG^*** is the binary OGG content matrix and has the same dimensions as ***D^OGG^***.

The phylogenetic profile of an OGG was defined as:

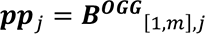

where ***pp****j* is the phylogenetic profile of OGG *j*, and *B*^*OGG*^_[1,*m*],*j*_ and *m* are the *j^th^* column vector and number of rows in the ***B^OGG^*** respectively. For each pair of proteins in the proteome of *E. coli* K12, a phylogenetic profiling feature of protein coevolution was defined as the average of the MI shared between the profiles of all pairs of OGGs encoded in the two proteins:

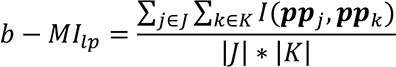

Where *b* − *MI*_*l,p*_ is the MI shared between the phylogenetic profiles of protein *l* and protein *p*, *J* and *K* are the sets of OGGs encoded in proteins *l* and *p* respectively, *j* and *k* are elements of *J* and *K* respectively, *pp*_*j*_ and *pp*_*k*_ are the phylogenetic profiles of *j* and *k* respectively, *Ipp*_*j*_, *pp*_*k*_) is the MI shared between *pp*_*j*_ and *pp*_*k*_ computed using Shannon’s classic formulation for the MI between two discrete random variables (Shannon, 1970; Cover and Thomas, 2006), and *|J|* and *|K|* are the number of elements in *J* and *K* respectively.

#### Covariation feature

The covariation between a pair of OGGs was described by:

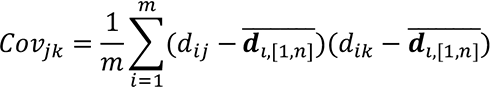

where *Cov*_*jk*_ is the covariation between OGGs *j* and *k*, *m* and *n* are the number of proteomes (rows) and OGGs (columns) in ***D^OGG^*** respectively, *dij* and *dik* are the number of annotations of OGGs *j* and *k* in proteome *i* respectively, and 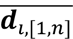 is the average number of OGG annotations in proteome *i* obtained by averaging the corresponding row vector in ***D^OGG^***. For each pair of proteins in the proteome of *E. coli* K12, protein covariation was defined as the average of the covariation shared between all pairs of OGGs encoded in the two proteins:

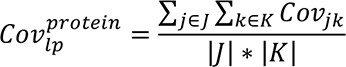

where 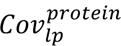 is the covariation feature of interaction between protein *l* and protein *p*, *J* and *K* are the sets OGGs encoded in proteins *l* and *p* respectively, *j* and *k* are elements of *J* and *K* respectively, *Cov*_*jk*_ is the covariation between OGGs *j* and *k*, and *|J|* and *|K|* are the number of elements in *J* and *K* respectively.

#### PCA-based spectral correlations features

These features were inspired by the approach of Franceschini and colleagues and the typical use of SVD to produce a low rank approximation of the initial data matrix in an effort to ‘denoise’ the data (Franceschini *et al*., 2016). For each pair of proteins in the *E. coli* K12 proteome spectral correlations were computed as described in the section ‘*Computing protein-protein spectral correlations’* over windows ranging from component 1 to component *k*, where *k* = 5, 10, 20, 50, 100, 500, 1000, 5000, or 7047.

### Validating RF models in two additional validation tasks

#### Training dataset task

Each decision tree within an RF model was tasked with predicting interaction classes for the out-of-bag examples from the training datasets. F-scores were computed for the consensus predictions of each model.

#### Comprehensive benchmark task

Biological interactions are typically sparse: the number of not-interacting pairs of proteins vastly outnumber the number of interacting pairs. As such, we desired to challenge each of the RF models in a validation task reflective of this asymmetry. To do so, each RF model was tasked with predicting classes for all pairs of proteins in the *E. coli* K12 proteome after exclusion of pairs used in the gold-standard dataset. These predictions were validated against four different comprehensive benchmarks: the indirect PPIs in the STRING Nonbinding benchmark (n = 5,423 indirect PPIs, 9,637,213 not-interacting), the mixed indirect/direct PPIs in the GO (n = 79,794 indirect or direct PPIs, 9,562,842 not-interacting) and STRING benchmarks (n = 20,216 indirect or direct PPIs, 9,622,420 not-interacting), and the direct PPIs in the entire PDB benchmark (n = 809 direct PPIs, 9,614,827 not-interacting).

### Predicting proteome-wide direct PPIs for 11 phylogenetically unrelated bacteria

#### Proteomes represented in **D**^OGG^

Each of the fifty RF models trained to classify interactions in *E. coli* K12 using MIWSCs were used to predict proteome-wide indirect and direct PPIs in the following bacteria (Uniprot Proteome ID, NCBI taxonomy ID in parentheses): *Aliivibrio fischeri* ES114 (UP000000537, 312309), *Azotobacter vinelandii* DJ (UP000002424, 322710), *Bacillus subtilis* 168 (UP000001570, 224308), *Caulobacter vibrioides* (UP000053705, 155892), *Helicobacter pylori* 26695 (UP000000429, 85962), *Mycobacterium tuberculosis* H37Rv (UP000001584, 83332), *Mycoplasma genitalium* G37 (UP000000807, 243273), *Pseudomonas fluorescens* F113 (UP000005437, 1114970), *Staphylococcus aureus* NCTC 8325 (UP000008816, 93061), *Streptomyces coelicolor* A3(2) (UP000001973, 100226), *Synechocystis sp.* PCC 6803 (UP000001425, 1111708). For each proteome, a set of consensus PPIs was defined as those for which a majority of the models (> 25) produced the same classification of ‘indirect PPI’ or ‘direct PPI’.

#### Proteomes not represented in **D**^OGG^

To predict interactions for a proteome that was not represented in ***D^OGG^*** (ex. *Azotobacter vinelandii* DJ, UP000002424, 322710), OGGs were mapped using EggNOG mapper V2 and MIWSCs were extracted using the OGG projections in ***V^OGG^*** (Huerta-Cepas, 2017; Huerta-Cepas, 2019). These features were used to predict proteome-wide indirect and direct PPIs as described for the Uniprot Reference Proteomes above.

### Validating direct PPI predictions against experimental evidence in the STRING database

The predicted direct PPIs were benchmarked against the sets of interactions in the STRING database with a non-zero experimental subchannel score for *E. coli* K12 and the eleven additional organisms described above.

### A head-to-head comparison with the approach of Cong and colleagues

Cong and colleagues have provided proteome-wide PPI predictions for *E. coli* K12 and *Mycobacterium tuberculosis* H37Rv (Cong *et al*., 2019). Their predictions of *E. coli* PPIs were based on amino acid coevolution supplemented with existing knowledge (‘Coev+’). In contrast, their predictions of PPIs in *M. tuberculosis* were based on amino acid coevolution alone (‘Coev’). Therefore, for a head-to-head comparison, we compared the predictions produced by our RF models trained on MIWSCs with their PPI predictions in *M. tuberculosis*. We benchmarked these interactions using two strategies. The first strategy mirrored that used by Cong and colleagues, computing the fraction of interactions assigned a STRING combined score of 0, 0-0.4, or > 0.4. The second strategy used orthogonal experimental evidence by computing the fraction of interactions assigned a STRING experimental subchannel score of 0 and > 0.

#### Generating **D**^domain^

PFAM domain annotations were downloaded from the UniProt database on 05/12/2020 (The UniProt Consortium, 2019). A domain count matrix was assembled (***D^domain^***, **Figure 3A**) where rows were defined as proteomes (n = 7,047, the same proteomes as described in **Figure 1 – figure supplement 1** and used to create ***D^OGG^***), columns were defined as domains, and the value in each cell was the number of annotations of a domain in a proteome. All domains present in fewer than 1% of the proteomes were removed leaving 7,245 unique columns in ***_D_domain***.

#### Performing SVD on **D**^domain^

SVD was performed on ***D^domain^*** following the same protocol described in the section ‘*Singular value decomposition (SVD) of D^OGG’^*. The SVD spectrum of ***D^domain^*** is displayed in **Figure 3B**, overlayed on the SVD spectrum derived from ***D^OGG^***.

### Relating the structure of protein covariation defined by domain-based features with the hierarchy of biological organization as shown in Figure 3C

#### Computing domain-based proteome-proteome and protein-protein spectral corelations

Domain-based proteome-proteome and protein-protein spectral correlations were computed in an identical fashion as described for OGG-based spectral correlations as described in the sections ‘*Computing proteome-proteome spectral correlations’* and ‘*Computing protein-protein spectral correlations’* respectively.

#### Computing MI shared between domain-based spectral correlations and benchmarks of biological organization

The MI shared between spectral correlations within all 5-component windows of the SVD spectrum of ***D^domain^***, ranging from component 1 to component 3000, and benchmarks of biological organization was computed in the identical way and for the identical benchmarks as described in the section *<u>Computing Mutual Information (MI) between spectral correlations and benchmarks of biological knowledge</u>* and is shown and compared to a random model in **Figure** 3 – figure supplement 1.

*Calculation of domain-based MI cdfs shown in* ***Figure 3C*** MI cdfs shown in **Figure 3C** were computed in an identical fashion as described in the section ‘*Calculation of MI cumulative distribution functions (cdfs) shown in* ***Figure 1C*.**

### Training and validating MIWSCdomain-based RF models to infer PPIs

RF models were trained using MIWSCdomains, validated, and compared to existing methods according to the same protocols described in the sections *Training and validating Random Forests (RF) models for predicting PPIs in E. coli K12 using MIWSCs*, *Training and validating*

*RF models on quantitative features of existing methods*, and *Validating RF models in two additional validation tasks*.

*Gene-set enrichment analysis performed on statistical model of E. coli K12 motility*

Gene-set enrichment analysis (GSEA) was performed on the sets of proteins defined by the statistical modules using DAVID analysis (v6.8). The ontological term with the lowest p-value is indicated for each statistical module shown in **Figure 4**. A full list of significant ontological terms and their associated p-values for each statistical module is listed in **Table S4**.

### Evaluating the robustness and generalizability of predicting the hierarchical organization of biological pathways using spectral correlations

For the additional analyses characterizing motility in *E. coli* K12 using MotB, characterizing motility in *B. subtilis* using Hag, characterizing amino acid metabolism in *E. coli* K12 using HisG, and characterizing motility in *E. coli* K12 and *B. subtilis* using FliC and Hag respectively with domain-based spectral correlations, the network graphs shown in **Figure 4 – figure supplement 3-6** were generated following the identical approach described in the section ‘*Predicting the hierarchical organization of the motility pathway in E. coli K12*’. Note that for the domain-based characterization, a separate null model for spectral correlations in *D^domain^* was developed and applied as described in the section *Developing a null model of random protein-protein spectral correlations within the SVD spectrum of **D******^OGG^***. Gene-set enrichment analysis was performed for each example of a statistically derived pathway in an identical fashion as described in the section *Gene-set enrichment analysis performed on statistical model of E. coli K12 motility*.

*Assaying strains of P. aeruginosa for pilus and flagellar motility*

All *P. aeruginosa* strains used in this study were ordered from the Manoil Lab. All strains were grown at 37°C on LB supplemented with 25µg/ml irgasan and gentamicin (75 µg/ml) as necessary. *E. coli* XL1-Blue was maintained on LB agar plates with gentamicin (15 µg/ml) as necessary.

*P. aeruginosa* transposon mutants of interest were ordered from the Manoil Lab. *P. aeruginosa* growth was at 37°C on LB supplemented with 25µg/ml irgasan and gentamicin (75 µg/ml) as necessary. Strains were assayed for subsurface twitching motility as previously described (Alm and Mattick, 1995; Little *et al*, 2018). Strains were grown overnight and stab inoculated in the interstitial space between the basal surface of 1.0% LB agar and a plastic petri dish. Plates were incubated for 48 hours at 37°C. Agar was removed and cells attached to the plate were stained with 0.5% crystal violet; twitch zone diameter was measured and plates were imaged.

Surface twitching motility assays were performed as previously described (Little *et al*., 2018; Kearns *et al*., 2001). *P. aeruginosa* strains of interest were grown overnight and concentrated in morpholinepropanesulfonic acid (MOPS) buffer (10mM MOPS, 8mM MgSO4, pH 7.6). A 2.5µl volume of the MOPS buffered bacterial suspension was spotted onto buffered twitching motility plates (10mM Tris, 8mM MgSO4, 1mM NaPO4, 0.5% glucose, 1.5% agar, pH 7.6) and was incubated for 24 hours at 37°C. The twitching zone was measured and imaged.

Swimming motility was performed as previously described (Rashid and Kornberg, 2000).

Overnight cultures were stab inoculated into the surface of LB-0.3% bacto agar and were incubated for 24 hours at 37°C. The resulting swimming zone was measured.

For complementation of genes of interest into *P. aeruginosa* strains, the complementation vector pBBR1-MCS5-PA0769 was created using Gibson assembly. The vector was transferred to *P. aeruginosa* by electroporation using 2.2kV in a 2mm gap cuvette, and subsequent selection using gentamicin.

## Tables with titles and Legends

Table S1: *D^OGG^* matrix, related to Figure 1A.

**Table S2: NCBI taxonomic strings for each organism used to generate phylogenetic benchmarks, related to Figure 1B.**

Table S3: Benchmarks of PPIs in *E. coli* K12, related to Figure 1B.

**Table S4: Data and statistical support for RF model validation studies, related to Figure 2, Figure 2—Figure Supplement 3.**

Table S5: *D^domain^* matrix, related to Figure 3.

**Table S6: Data and statistical support for domain-based RF model validation studies, related to Figure 3—Figure Supplement 2.**

**Table S7: Data pertaining to Figure 4.**

**Table S8: Data related to Figure 4—Figure Supplement 3.**

**Table S9: Data related to Figure 4—Figure Supplement 4.**

**Table S10: Data related to Figure 4—Figure Supplement 5.**

**Table S11: Data related to Figure 4—Figure Supplement 6.**

**Table S12: Data related to Figure 4—Figure Supplement 6.**

**Table S13: Data related to Figure 5A,B and Figure 5—Figure Supplement 1.**

**Table S14: Data related to Figure 5C.**

**All Tables can be downloaded at github.com/arjunsraman/Zaydman_et_al as .zip files (Tables_S1_to_S4.zip, Tables_S5_to_S14.zip).**

## Acknowledgments

We thank Robert Y. Chen, Adam Bailey, Nima Mosammaparast, and Jacqueline Payton for substantial discussion regarding this manuscript. We thank Rama Ranganathan for a critical reading of the manuscript as well as in-depth discussion. We thank Dinanath Sulakhe (Center for Research Informatics (CRI), University of Chicago) for assisting in producing the web application tools described in this manuscript. We thank Sam Light, Sampriti Mukherjee, and Eric Pamer for helpful discussions regarding experiments performed.

## Author Contributions

M.A.Z. and A.S.R. conceived the project, developed the mathematical approaches described, wrote the code for conducting analyses, performed data analysis, and assembled the manuscript. A.L., performed the experiments regarding *P. aeruginosa*. W.J.B. provided technical expertise and wrote code for annotation and parsing and contributed to the writing of the manuscript. A.D., J.I.G., and J.M. contributed to this manuscript by providing critical feedback, in-depth discussion, and contributing to the writing of this manuscript.

## Figure Supplements

**Figure 1 – figure supplement 1.**
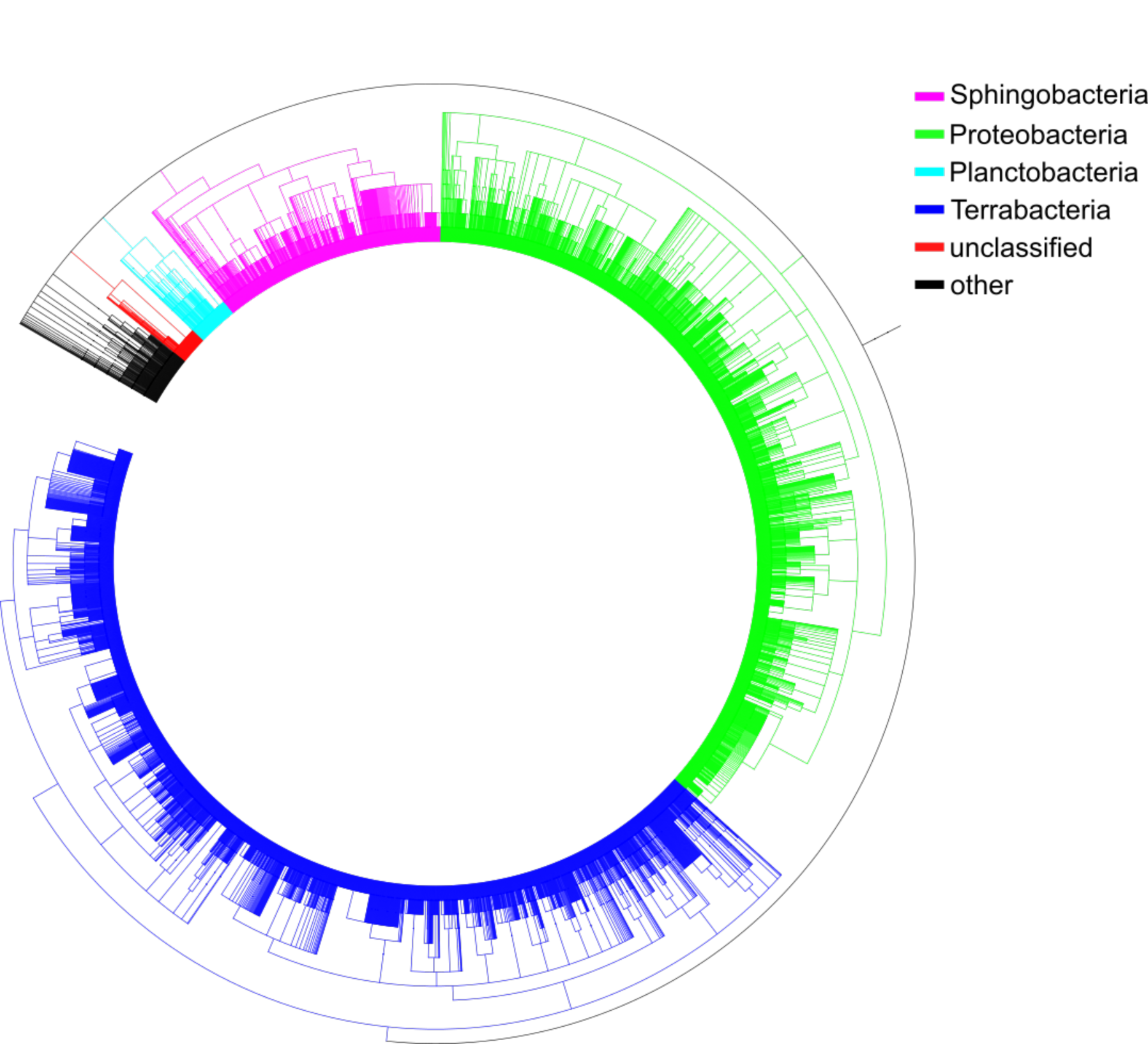
Phylogenetic tree of the 7,047 bacterial species represented in the UniProt reference proteomes database (release 02/2020) that served as the substrate for ***D^OGG^*** (Figure 1A). The tree was generated using PhyloT based on the NCBI taxonomy database and visualized using iTOL (Letunic et al., 2006).

**Figure 1 – figure supplement 2.**
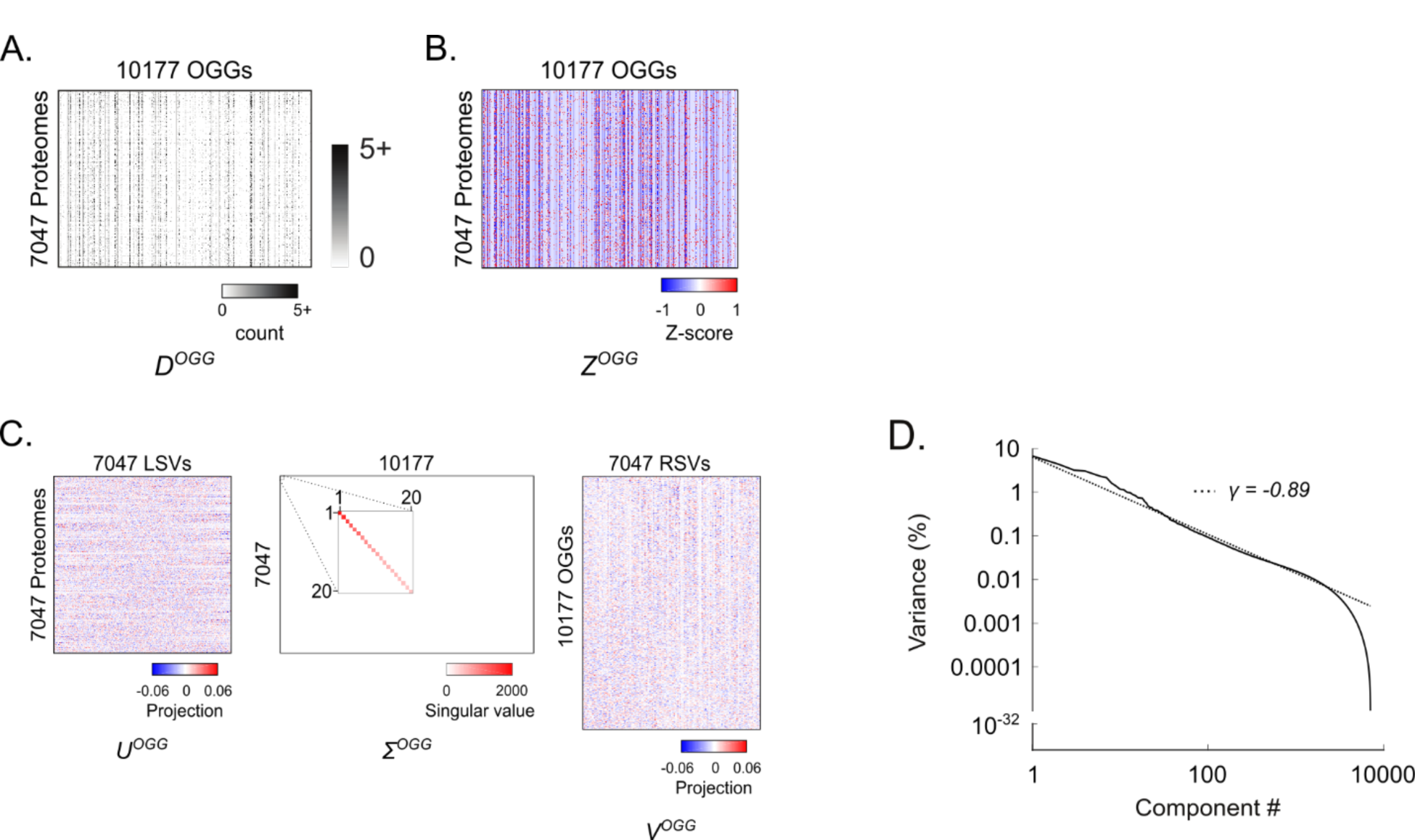
Using SVD to spectrally decompose OGG covariation in bacteria. (**A**) The raw OGG content matrix, ***D^OGG^***. (**B**) The z-scored OGG content matrix, ***Z^OGG^*** produced by subtracting the column mean from each element in ***D^OGG^*** and then dividing by the column standard deviation. (**C**) Application of SVD to ***Z^OGG^*** produced three matrices: *U^OGG^*, Σ*^OGG^*, and *V^OGG^*. *U^OGG^* contains the left singular vectors (LSVs) describing the projections of each proteome onto the SVD components. Σ*^OGG^* is a diagonal matrix with decreasing diagonal elements containing the singular values (inset expands top 20 singular values). The magnitude of each singular value is related to the fraction of the overall data variance explained by the corresponding SVD component. *V^OGG^* contains the right singular vectors (RSVs) describing the projections of each OGG onto the SVD components. (**D**) Percent variance explained per component versus component number. Dotted line represents the fit to a power-law distribution with the indicated exponent (*γ*).

**Figure 1 – figure supplement 3.**
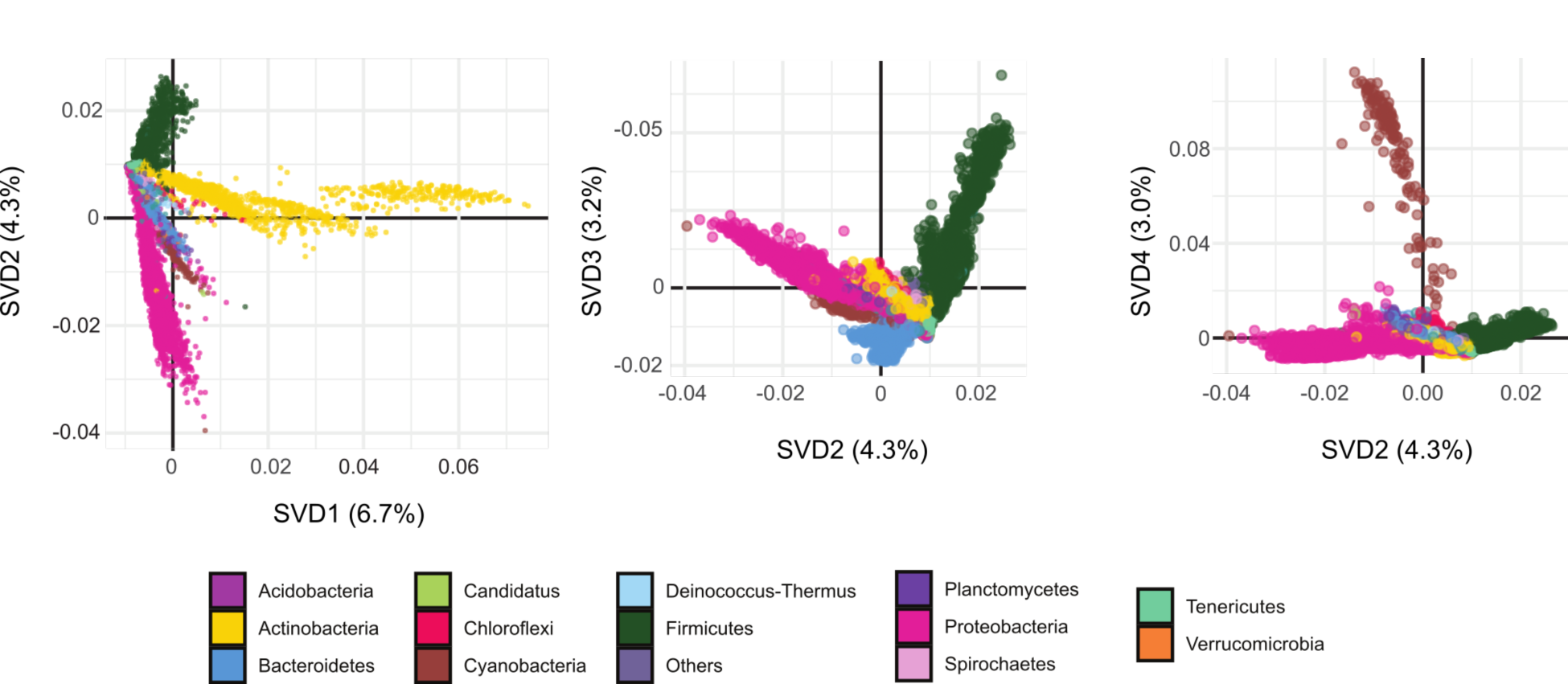
Projection of proteomes onto SVD1, SVD2, SVD3, and SVD4 of *D^OGG^* colored by phylum. Proteome projections onto the SVD components are derived from ***U^OGG^***, the matrix of left singular vectors (LSVs) defined by applying SVD to ***D^OGG^*** (Figure 1 **– figure supplement 2C**). Each marker represents a single proteome and is colored according by Phylum as indicated. The percent variance explained per SVD component is indicated in parentheses.

**Figure 1 – figure supplement 4.**
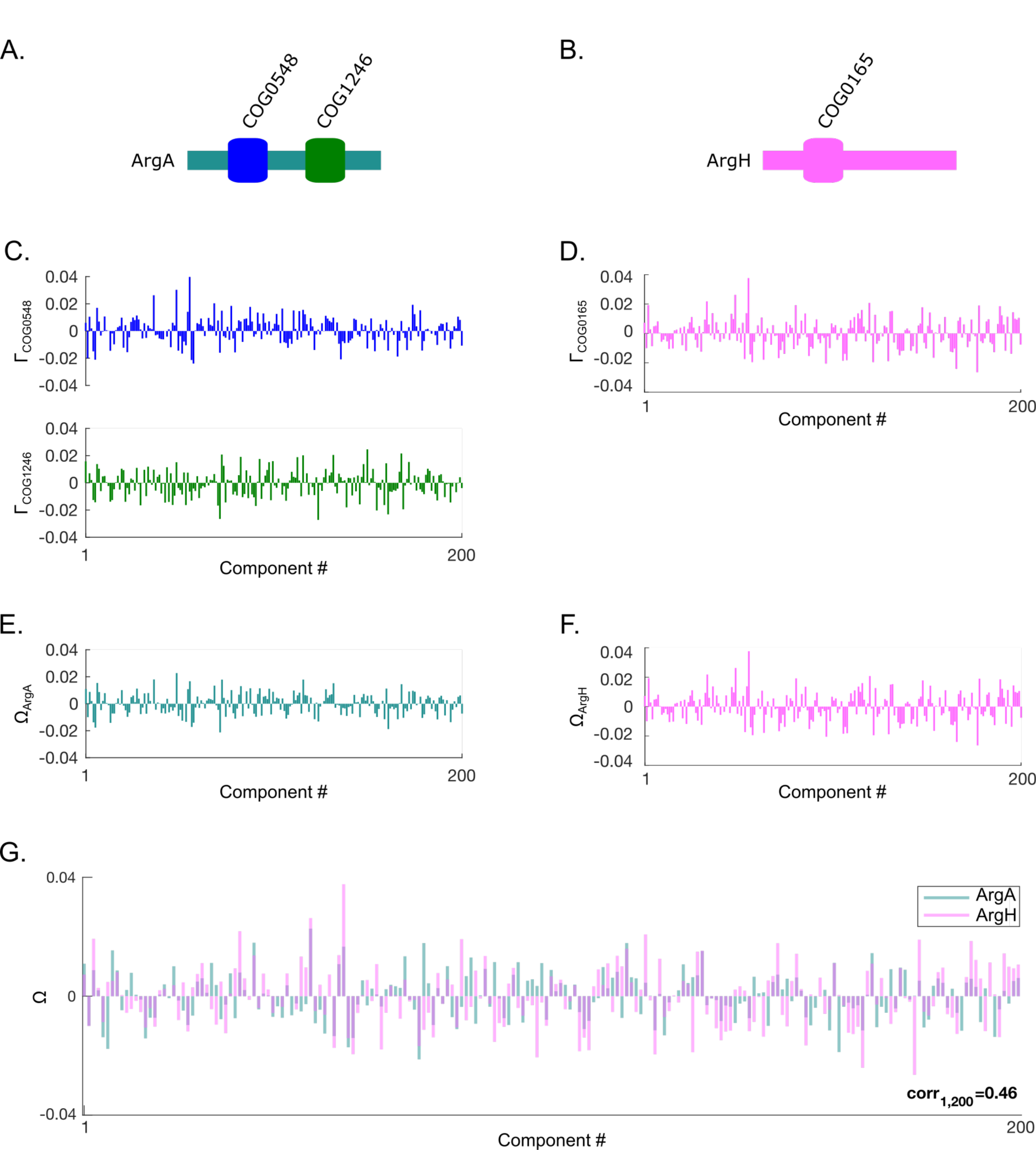
The process of computing spectral correlations between two proteins within a specific region of the SVD spectrum of *D^OGG^*. (**A,B**) The OGG structures of *E. coli* K12 ArgA (A) and ArgH (B). (**C,D**) The projections (Γ) of the OGGs encoded in ArgA (COG0548 and COG1246) (C) and ArgH (COG0165) (D) onto SVD1 to SVD200 of the SVD spectrum of *D^OGG^*. (**E,F**) The approximated projections (Ω) of ArgA (E) and ArgH (F) derived by averaging the projections for the OGGs encoded within each protein. (**G**) Overlay of the approximated protein projections of ArgA and ArgH. These two protein project similarly across SVD1 to SVD200 resulting in a positive spectral correlation value (Pearson correlation value shown).

**Figure 1 – figure supplement 5.**
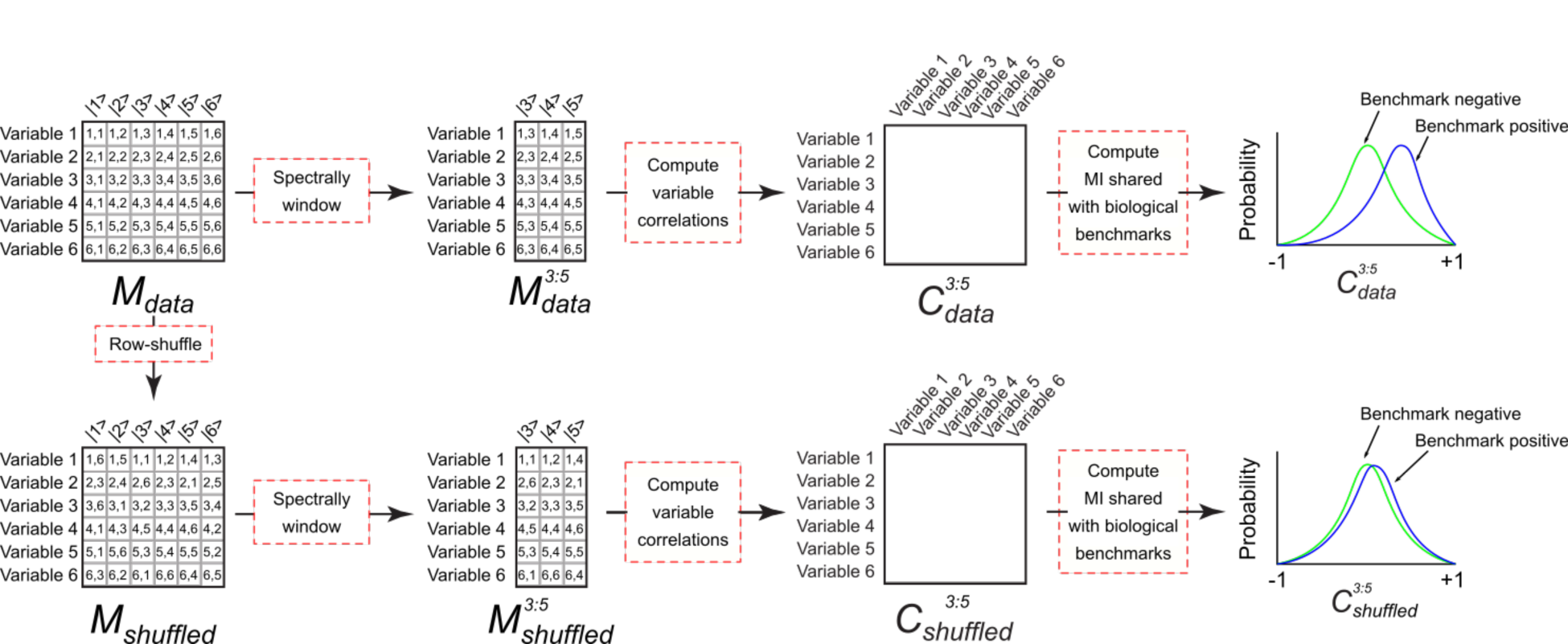
Estimating the MI shared between spectral correlations and a benchmark. (Top) Hypothetical projection matrix ***Mdata*** consists of projections of six variables onto six SVD components. If the variables correspond to the rows or columns of ***D***, the initial data matrix, then the matrix ***Mdata*** corresponds to the complete ***U*** or ***V*** matrices produced by application of SVD to ***D***, respectively. ***Mdata*** is windowed to components 3 to 5 to produce ***Mdata^3:5^*** by discarding all columns outside of this range. Next Pearson correlations are computed between all pairs of rows in ***Mdata^3:5^*** to produce the windowed spectral correlation matrix ***Cdata^3:5^***. The MI shared between ***Cdata^3:5^*** and a benchmark reflects the degree to which the distribution of spectral correlation values in ***Cdata^3:5^*** differs for variable pairs that interact within the benchmark (benchmark positive) and variable pairs that do not interact in the benchmark (benchmark negative). (Bottom) To estimate the MI produced by spurious correlations, ***Mdata*** is subjected to random row permutation to generate ***Mshuffled***. This process maintains the distribution of projection values for each variable but erases biologically meaningful spectral correlations leaving only the spurious correlations. ***Mshuffled*** is subjected to the identical windowing, row correlation computation, and MI calculations as described above for ***Mdata***. The biologically meaningful MI is estimated to be the difference between the MI estimate for ***Cdata^3:5^*** and ***3:5 Cshuffled*** .

**Figure 1 – figure supplement 6.**
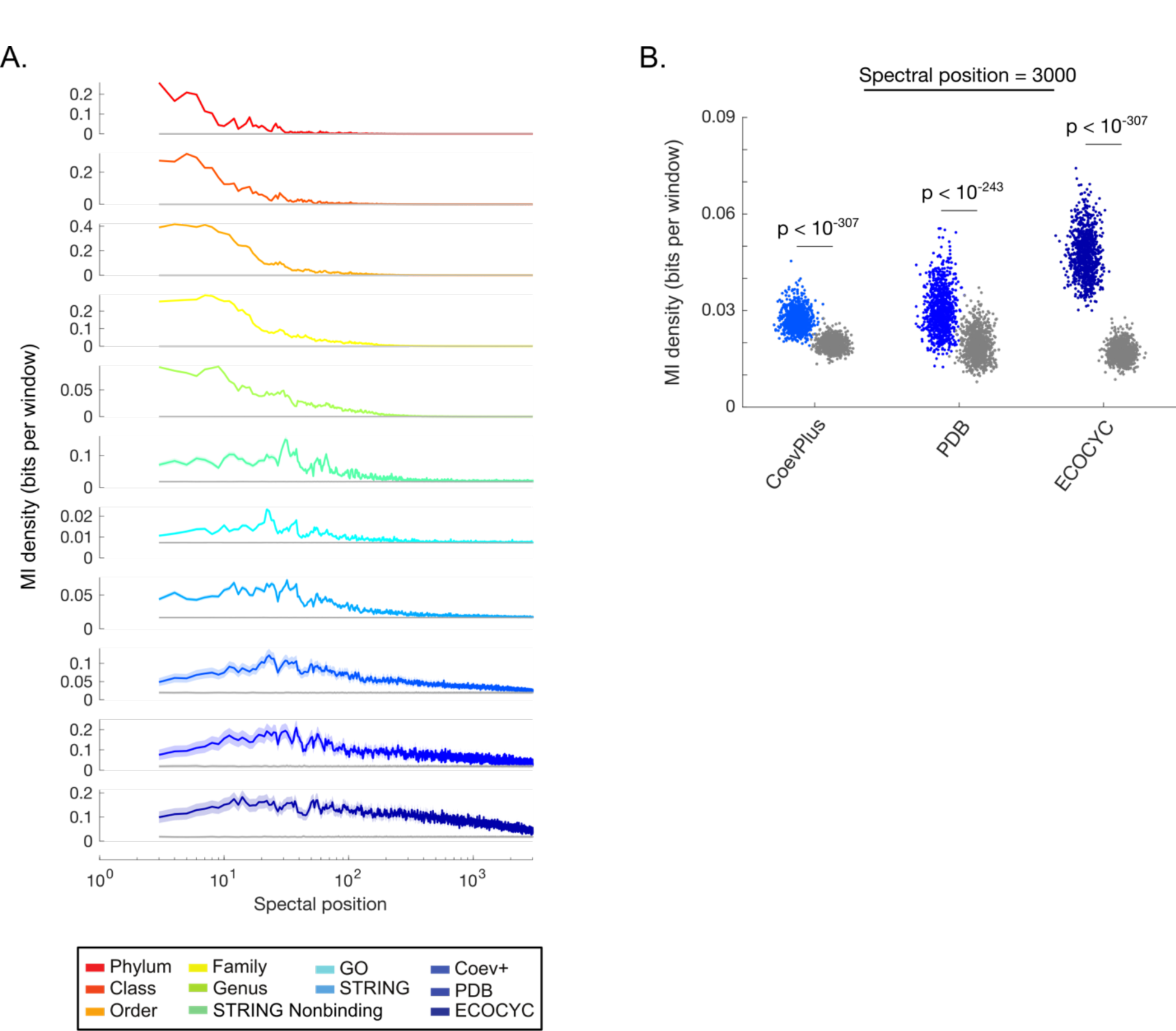
Quantifying the biological information contained within different regions of the SVD spectrum of *D^OGG^*. (**A**) MI density contained within a 5-component window versus spectral position for all windows between SVD1 and SVD3000 in the SVD spectrum of ***D^OGG^***. Colored and gray lines represent the MI values for the data matrix or the model of spurious spectral correlations, respectively. Lines and shaded contours represent the mean ± 2 standard deviations for the bootstraps of each benchmark. (**B**) MI density contained within SVD2995-SVD3000 of the SVD spectrum of ***D^OGG^***. Each dot represents the MI value for a single bootstrap of the indicated benchmark. Colored dots represent the MI for the data matrix and gray dots represent the MI for the model of spurious correlations. P-values produced by a paired Student’s t-test are indicated.

**Figure 2 – figure supplement 1.**
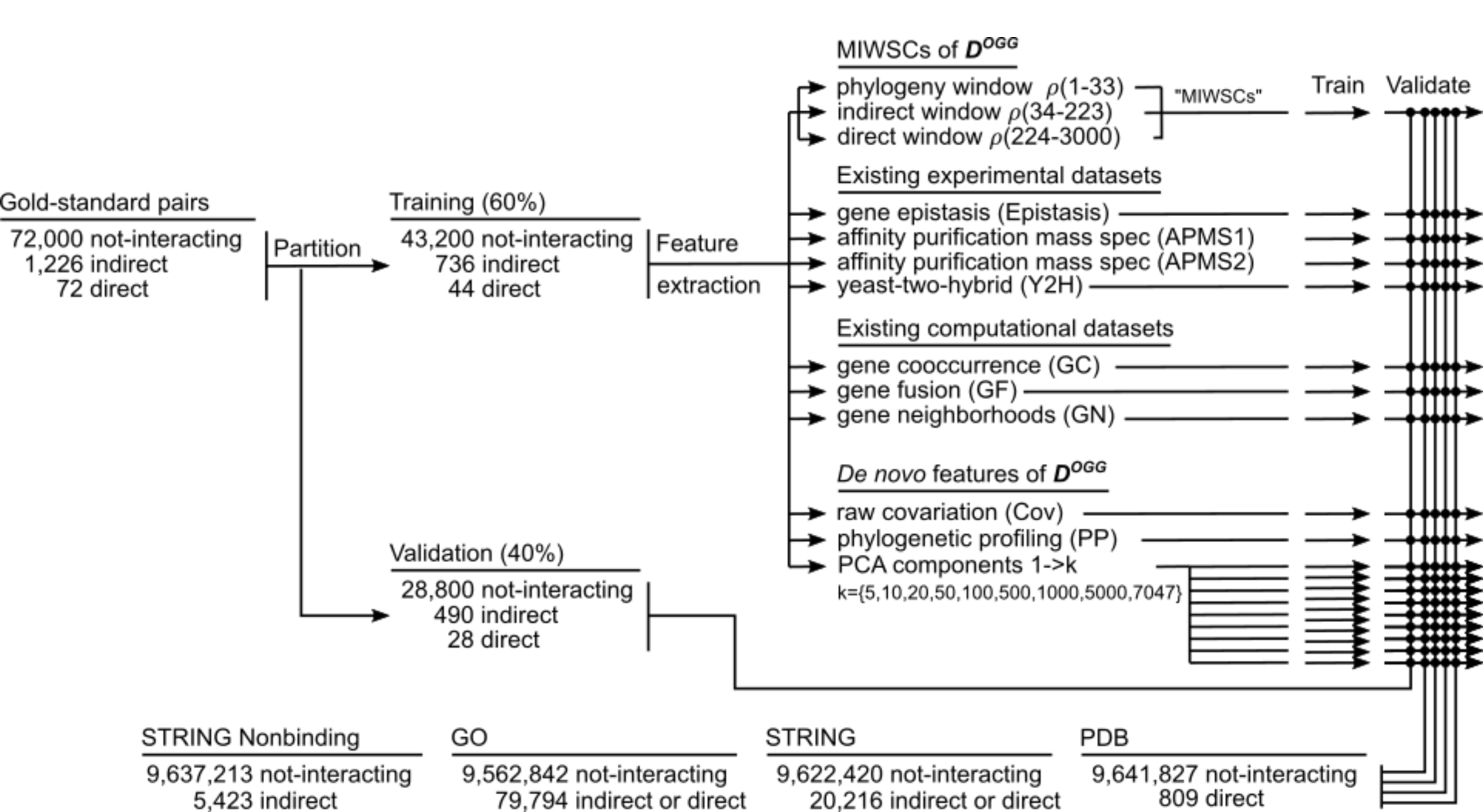
Multi-level classification task where RF models were trained and validated for predicting indirect and direct PPIs in *E. coli* K12 using spectral correlations features, existing computational features, or existing experimental features. A gold-standard dataset of well-characterized *E. coli* K12 protein pairs was assembled and partitioned into training and validation datasets. The labeled examples from the training set were used to train RF models to classify protein interactions using different features as indicated. The performance of the various RF models was benchmarked and compared by computing F-scores for classifying PPIs in the validation dataset, out-of-bag examples in the training dataset, and PPIs in four additional comprehensive benchmarks. This process of partitioning the gold-standard dataset, training, and validation was repeated 50 times to evaluate the reproducibility of RF model performance.

**Figure 2 – figure supplement 2.**
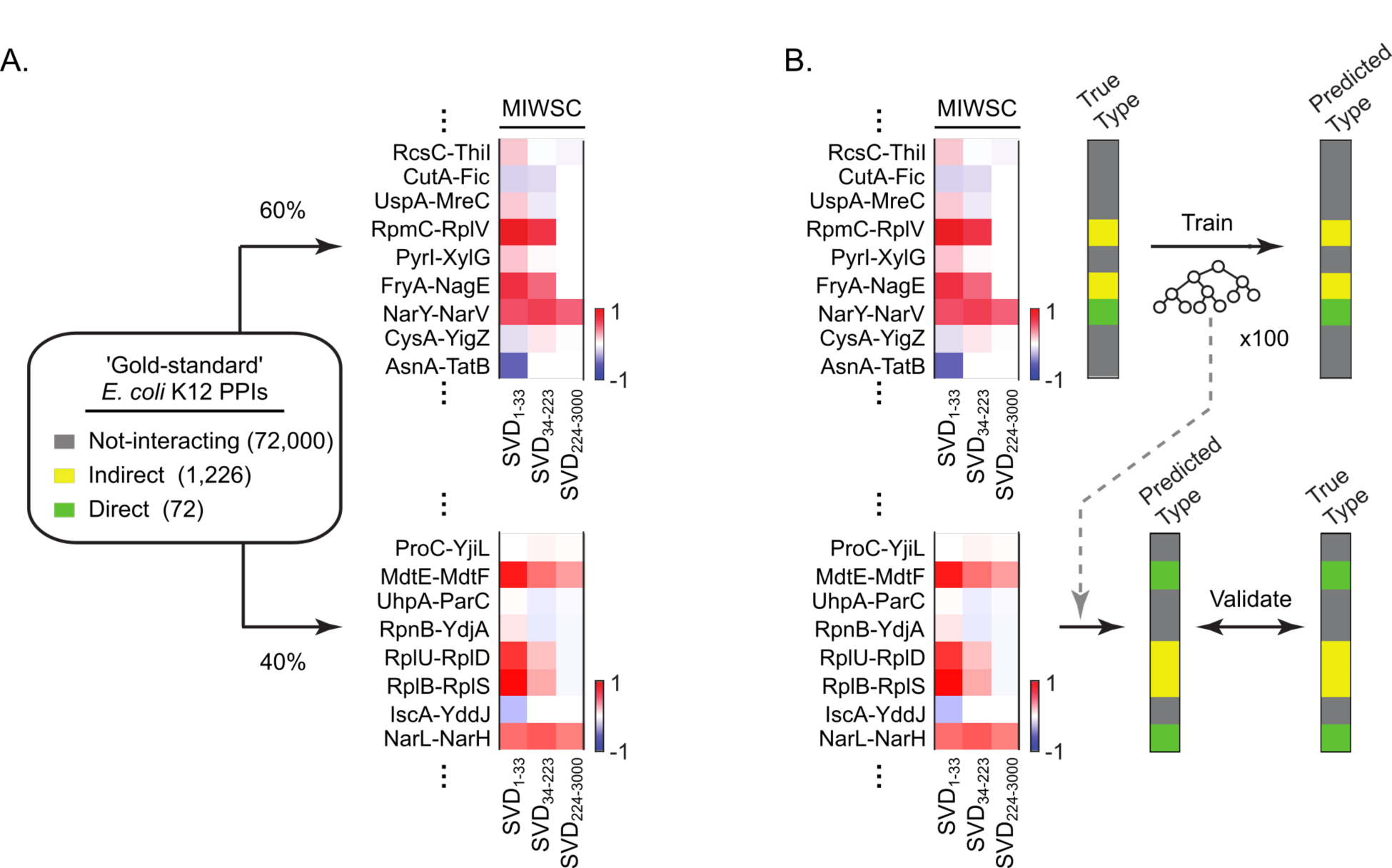
Extracting MIWSC features for the training and validation of RF models to classify *E. coli* K12 protein pairs as not-interacting (‘not-interacting’), indirect PPI (‘indirect’), or direct PPI (‘direct’). (A) A gold standard dataset of well characterized *E. coli* K12 protein pairs was assembled and randomly partitioned in to training (60%) and validation (40%) datasets. An MIWSC feature was extracted for each protein pair in the training and validation partitions of the gold standard dataset. The MIWSC feature consists of a set of three spectral correlations computed across SVD1-33, SVD34-223, and SVD234-3000. Each pixel of the heat map is the spectral correlation for the protein pair indicated in the row across the SVD components indicated in the column. (B) Using only the MIWSC features, RF models were trained using the labeled examples in the training dataset and then challenged to predict the unlabeled examples in the validation dataset.

**Figure 2 – figure supplement 3.**
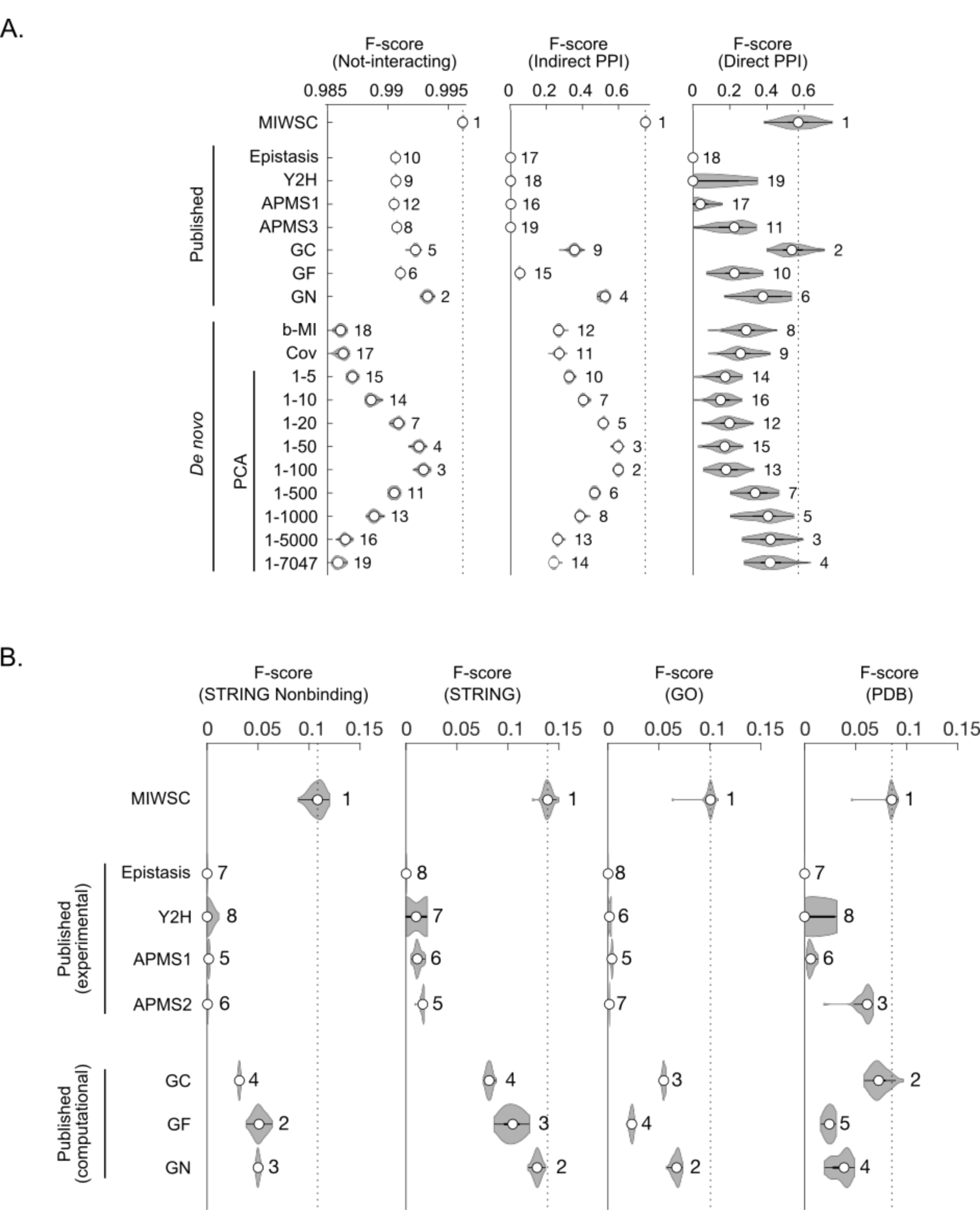
F-scores for predicting interaction classes for out-of-bag examples in the training datasets (A) and four additional comprehensive benchmarks (B). The violin plots describe the distribution of F-scores for models trained and validated on 50 random partitions of the gold-standard dataset (Figure 2 **– figure supplement 1**). Numbering indicates the rank of the median F-score for models trained on each feature (**Table S4**). Feature descriptions can be found in the legend of Figure 2.

**Figure 3 – figure supplement 1.**
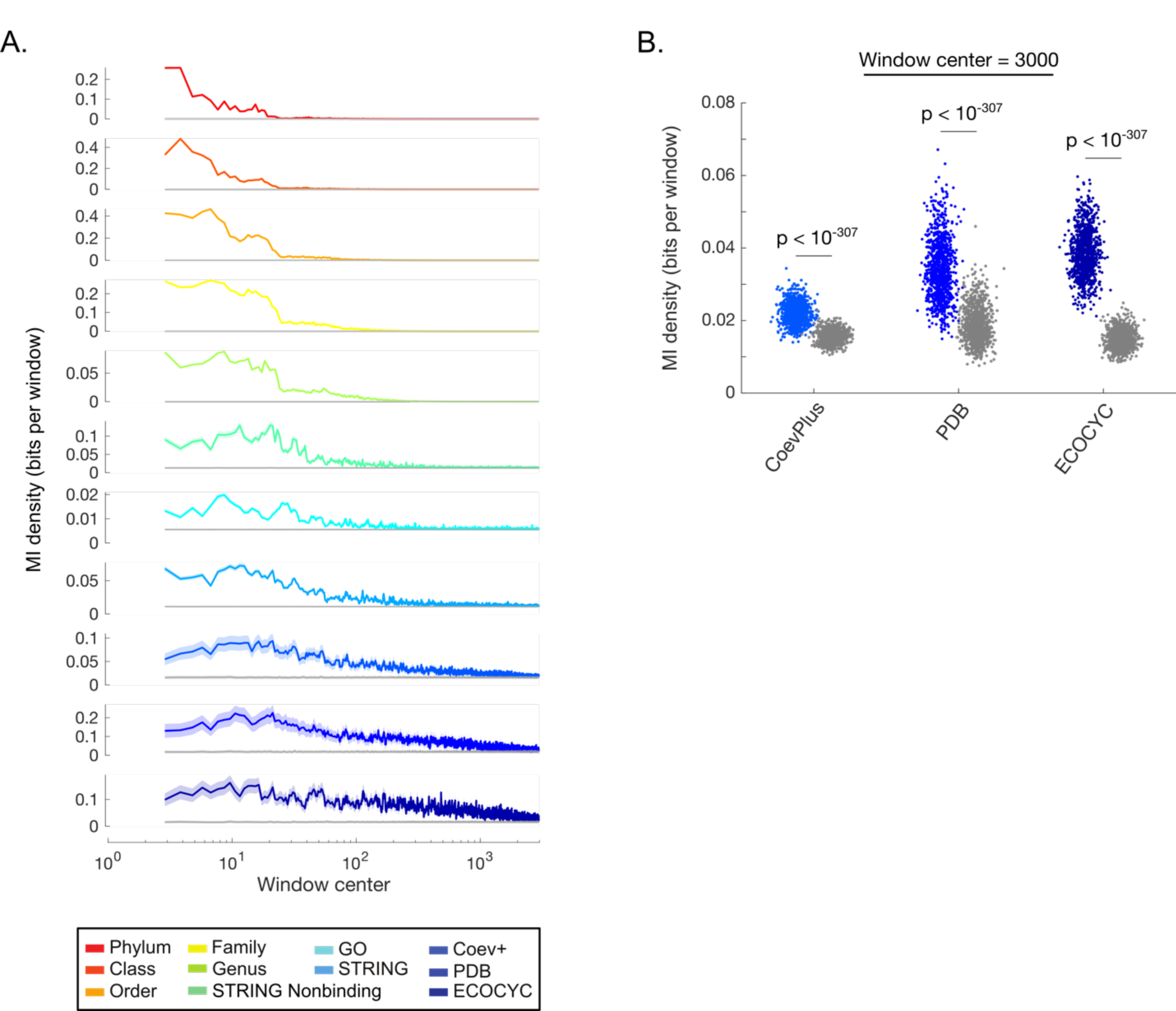
Quantifying the biological information contained within different regions of the SVD spectrum of *D^domain^*. (**A**) MI density contained within a 5-component window versus spectral position for all windows between SVD1 and SVD3000 in the SVD spectrum of ***D^domain^***. Colored and gray lines represent the MI values for the data matrix or the model of spurious spectral correlations, respectively. Lines and shaded contours represent the mean ± 2 standard deviations for the bootstraps of each benchmark. (**B**) MI density contained within SVD2995-SVD3000 of the SVD spectrum of ***D^domain^***. Each dot represents the MI value for a single bootstrap of the indicated benchmark. Colored dots represent the MI for the data matrix and gray dots represent the MI for the model of spurious correlations. P-values produced by a paired Student’s t-test are indicated.

**Figure 3 – figure supplement 2.**
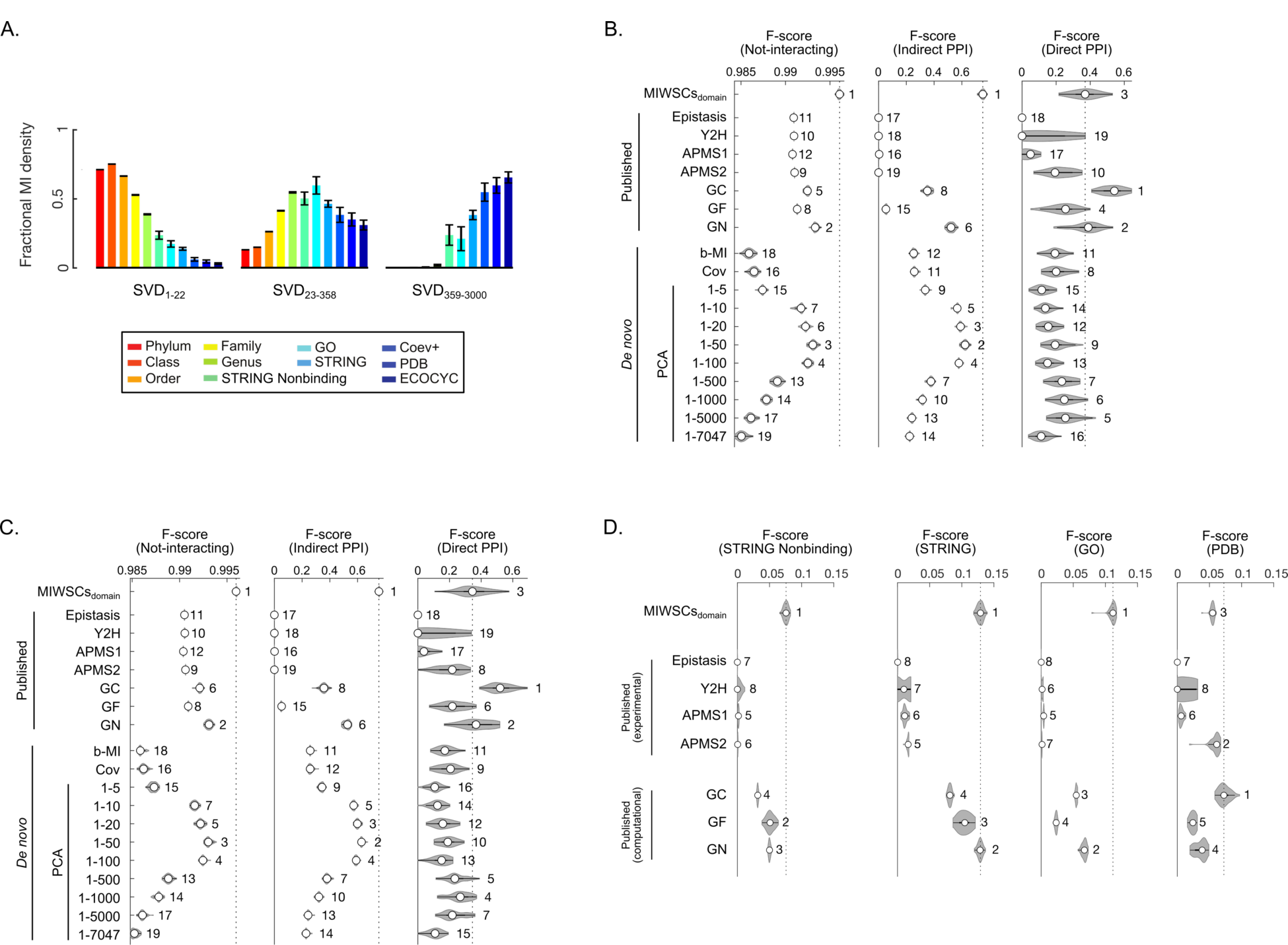
Domain-based MI windowed spectral correlations (MIWSCsdomain) enable accurate classification of protein pairs as either not-interacting, indirect PPI, or direct PPI. (A) Fractional MI density regarding each benchmark contained within spectral correlations computed across SVD1-22, SVD23-358, or SVD359-3000 of *D^domain^*. Color scheme is defined in the legend and follows that of Figure 1B**,C**. (**B-D**) F-scores for classifying protein pairs in the validation dataset (**B**), out-of-bag examples from the training dataset (**C**), or additional examples in four comprehensive benchmarks (**D**). The violin plots describe the distribution of F-scores for models trained and validated on 50 random partitions of the gold-standard dataset (Figure 2 **– figure supplement 1**). Numbering indicates the rank of the median F-score for models trained on each feature (**Table S6**). Feature descriptions can be found in the legend of Figure 2.

**Figure 4 – figure supplement 1.**
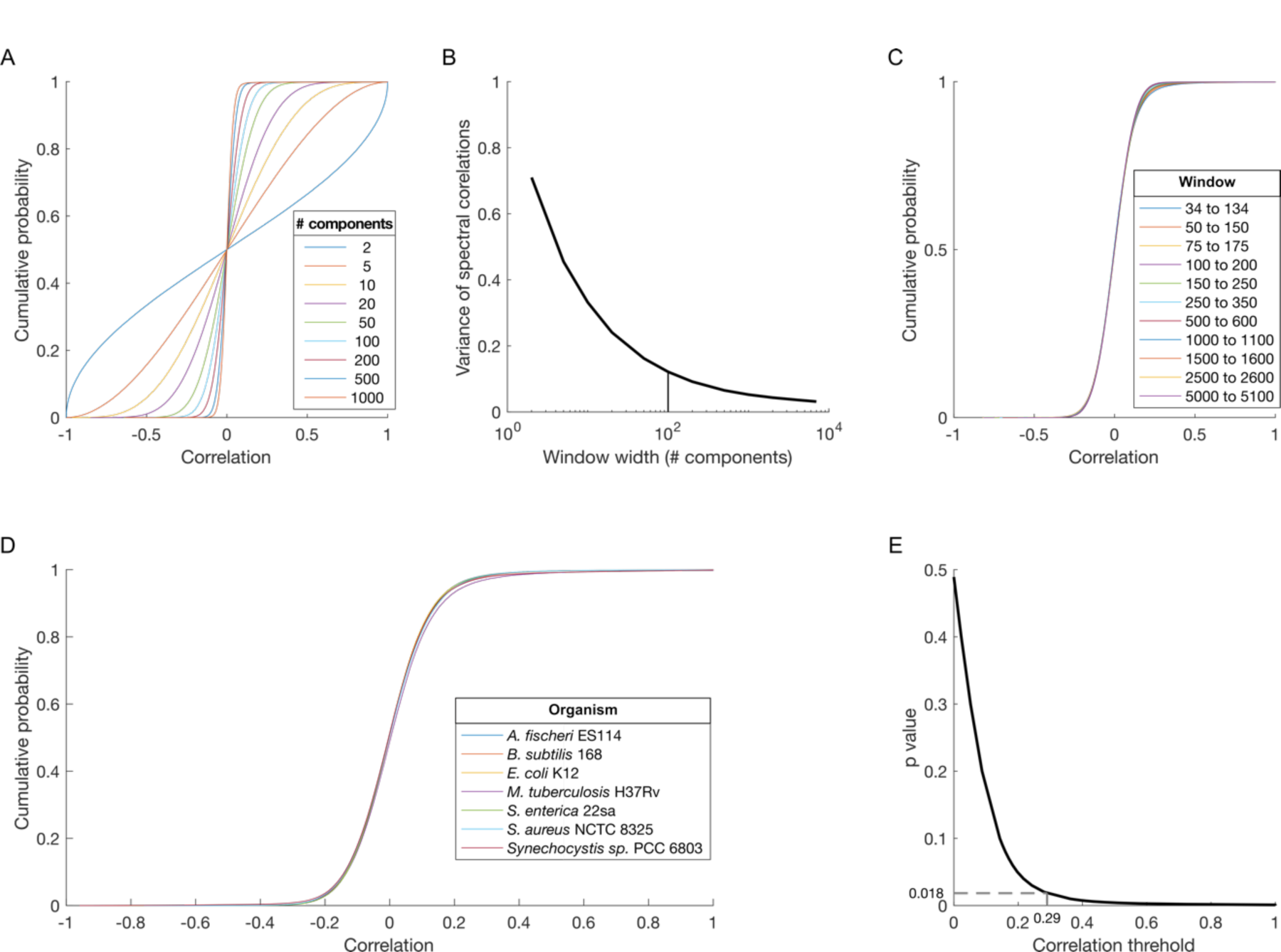
Developing a null model of random protein-protein spectral correlations within the SVD spectrum of *D^OGG^*. (**A**) Cumulative distribution functions (cdfs) for spectral correlations between all proteins in *E. coli* K12 across windows of different widths (legend) centered on component 1001. (**B**) Variance of the distributions in panel A plotted versus window width. Vertical line indicates a window width of 100 components. (**C**) cdfs for spectral correlations between all proteins in *E. coli* K12 across the indicated 100-component spectral windows (legend). (**D**) cdfs for spectral correlations for all proteins in proteomes from diverse organisms (legend) across the 100-component window centered on component 84. (**E**) p-value versus correlation threshold for spectral correlations between proteins in *E. coli* K12 across SVD34 to SVD134. The chosen correlation threshold (0.29) and associated p-value (0.018) are indicated.

**Figure 4 – figure supplement 2.**
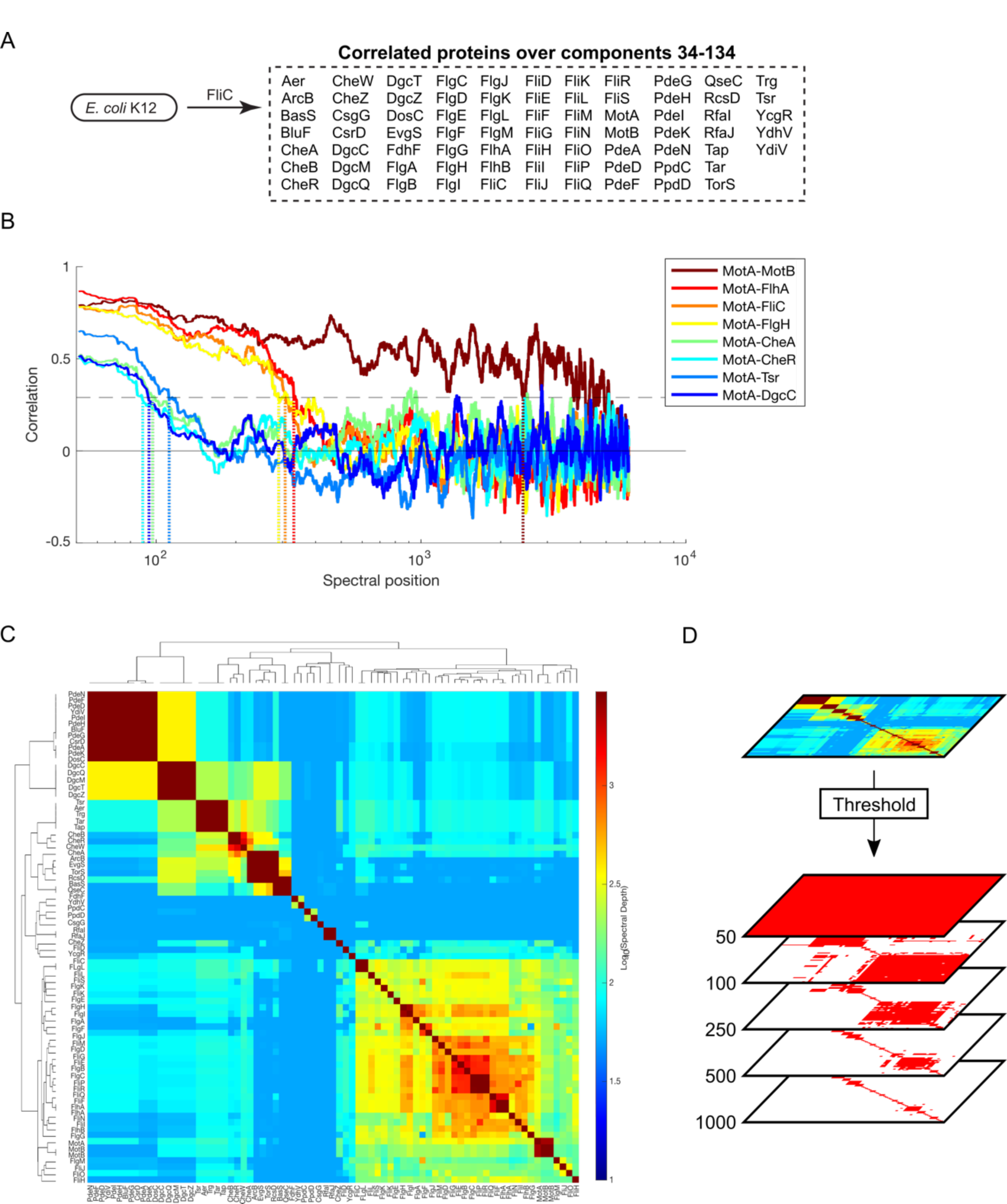
Process for constructing the FliC interaction networks appearing in Figure 4 using thresholded spectral depth. **(A)** Proteins that shared significant spectral correlations with FliC across SVD34 to SVD134. **(B)** Pairwise spectral correlations between indicated protein pairs (legend) as a function of spectral position. Horizontal dashed line represents the threshold of significant spectral correlation described in Figure 4 **– figure supplement 1E**. Dashed vertical lines represent the ‘spectral depth’—the spectral position at which spectral correlation first falls below the significance threshold. **(C)** Hierarchically clustered spectral depth matrix for all pairs of proteins in panel A. **(D)** Set of adjacency matrices derived from thresholding the spectral depth matrix in panel C. Red and white pixels indicate proteins that do or do not share a spectral depth exceeding the indicated thresholds respectively.

**Figure 4 – figure supplement 3.**
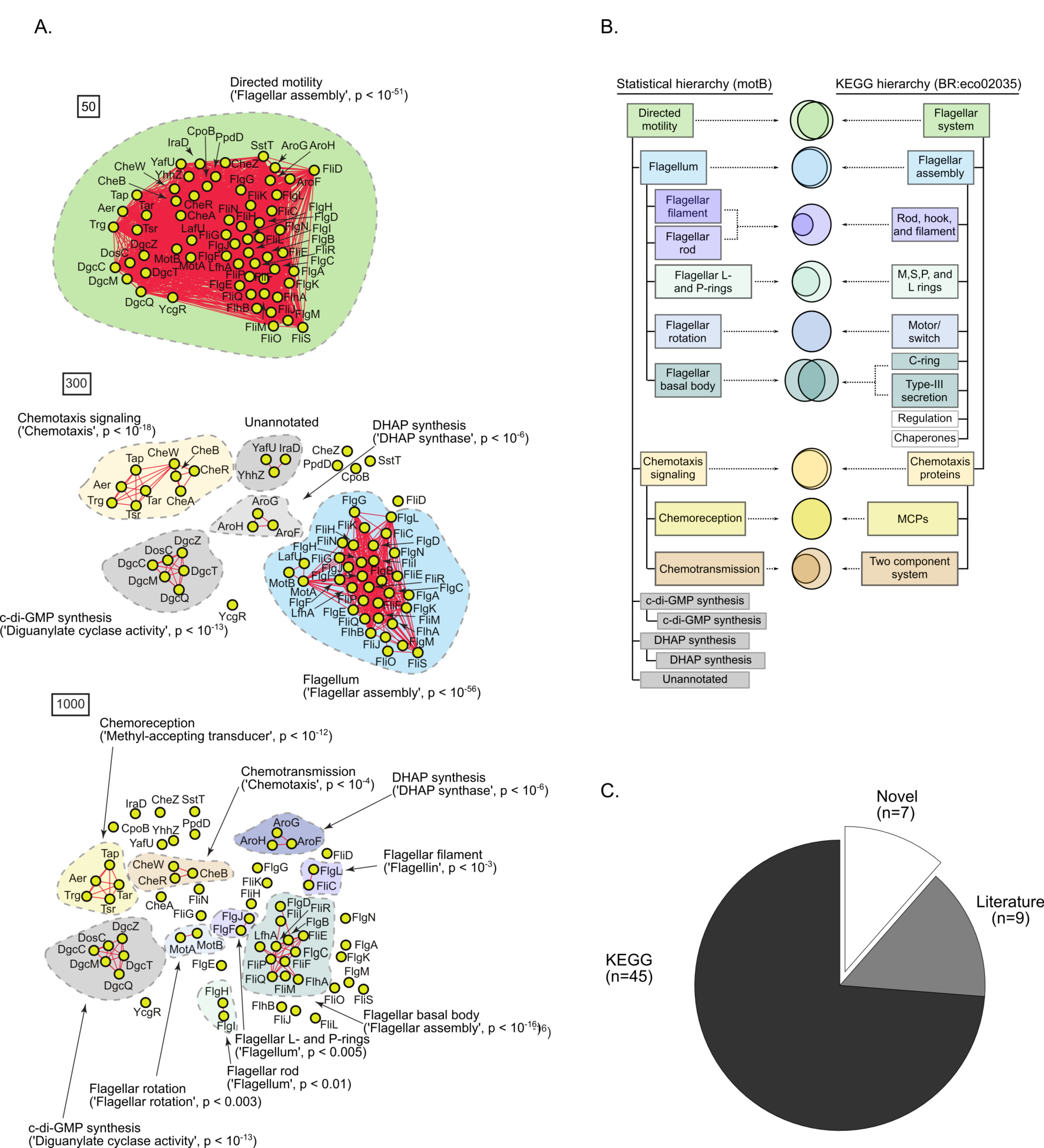
A statistically-derived hierarchical model of motility in *E. coli* K12 using MotB as a query protein. (**A**) 60 proteins in *E. coli* K12 shared significant spectral correlations with MotB across SVD34 to SVD134. Statistical interaction networks were defined by thresholding spectral depth at 50 (top panel), 300 (middle panel), and 1000 (bottom panel). Nodes (yellow circles) represent proteins and edges (red lines) represent statistical interactions. Shaded contours identify discrete subnetworks and are labeled with their assigned function based on interpretation of gene-set enrichment analysis (GSEA) and literature review. The most significantly enriched ontological term produced by GSEA and the associated p-value is shown in parentheses for each subnetwork (**Table S8**). B) Comparison of the statistically-derived model using MotB (left) to the KEGG model (BR:eco02035, right) of *E. coli* K12 motility. Venn diagrams represent the overlap between the sets of proteins in the indicated subnetwork of panel A (left) and the indicated KEGG category (right). (**C**) Pie graph of the number of proteins in the statistical model that are represented in the KEGG hierarchy (‘KEGG’), missing from the KEGG hierarchy but supported by experimental evidence in the literature (‘Literature’), or absent from the KEGG hierarchy and the literature (‘Novel’).

**Figure 4 – figure supplement 4.**
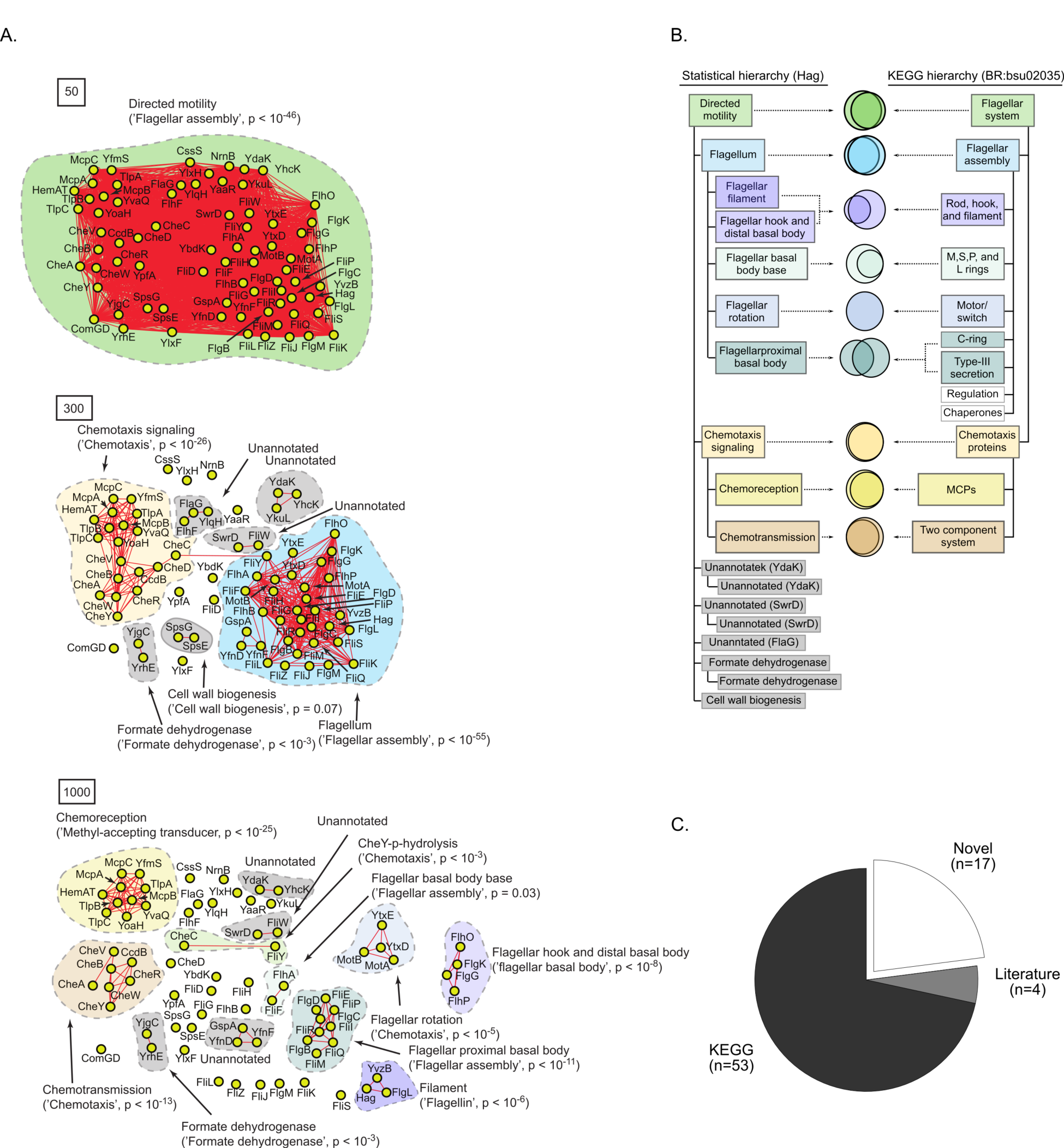
A statistically-derived hierarchical model of motility in *B. subtilis* 168 using Hag as a query protein. (**A**) 74 proteins in *B. subtilis* 168 shared significant spectral correlations with Hag over SVD34 to SVD134. Statistical interaction networks were defined by thresholding spectral depth at 50 (top panel), 300 (middle panel), and 1000 (bottom panel). Nodes (yellow circles) represent proteins and edges (red lines) represent statistical interactions. Shaded contours identify discrete subnetworks and are labeled with their assigned function based on interpretation of gene-set enrichment analysis (GSEA) and literature review. The most significantly enriched ontological term produced by GSEA and the associated p-value is shown in parentheses for each subnetwork (**Table S9**). B) Comparison of the statistically-derived model using Hag (left) to the KEGG model (BR:bsu02035, right) of *B. subtilis* 168 motility. Venn diagrams represent the overlap between the sets of proteins in the indicated subnetwork of panel A (left) and the indicated KEGG category (right). (**C**) Pie graph of the number of proteins in the statistical model that are represented in the KEGG hierarchy (‘KEGG’), missing from the KEGG hierarchy but supported by experimental evidence in the literature (‘Literature’), or absent from the KEGG hierarchy and the literature (‘Novel’).

**Figure 4 – figure supplement 5.**
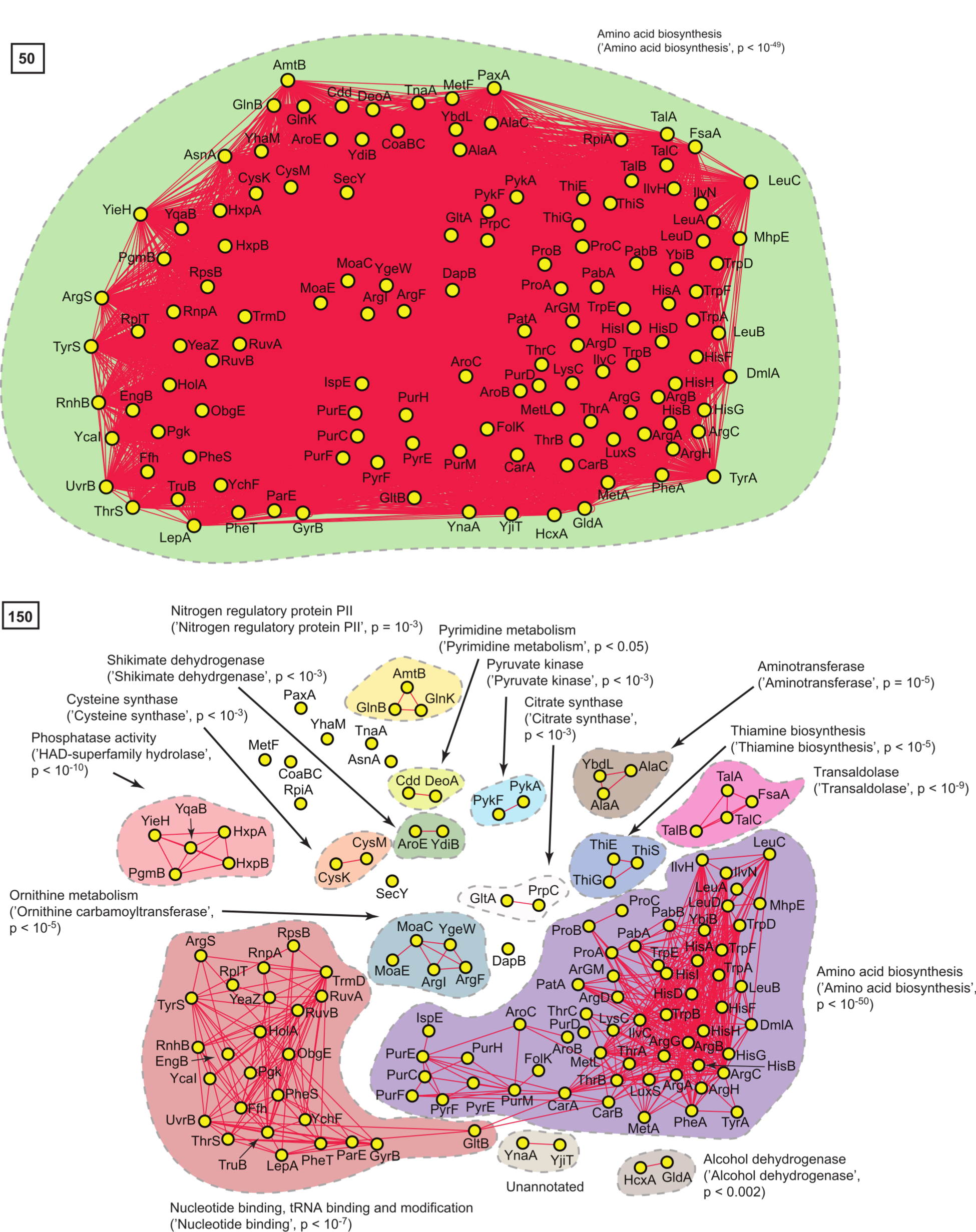
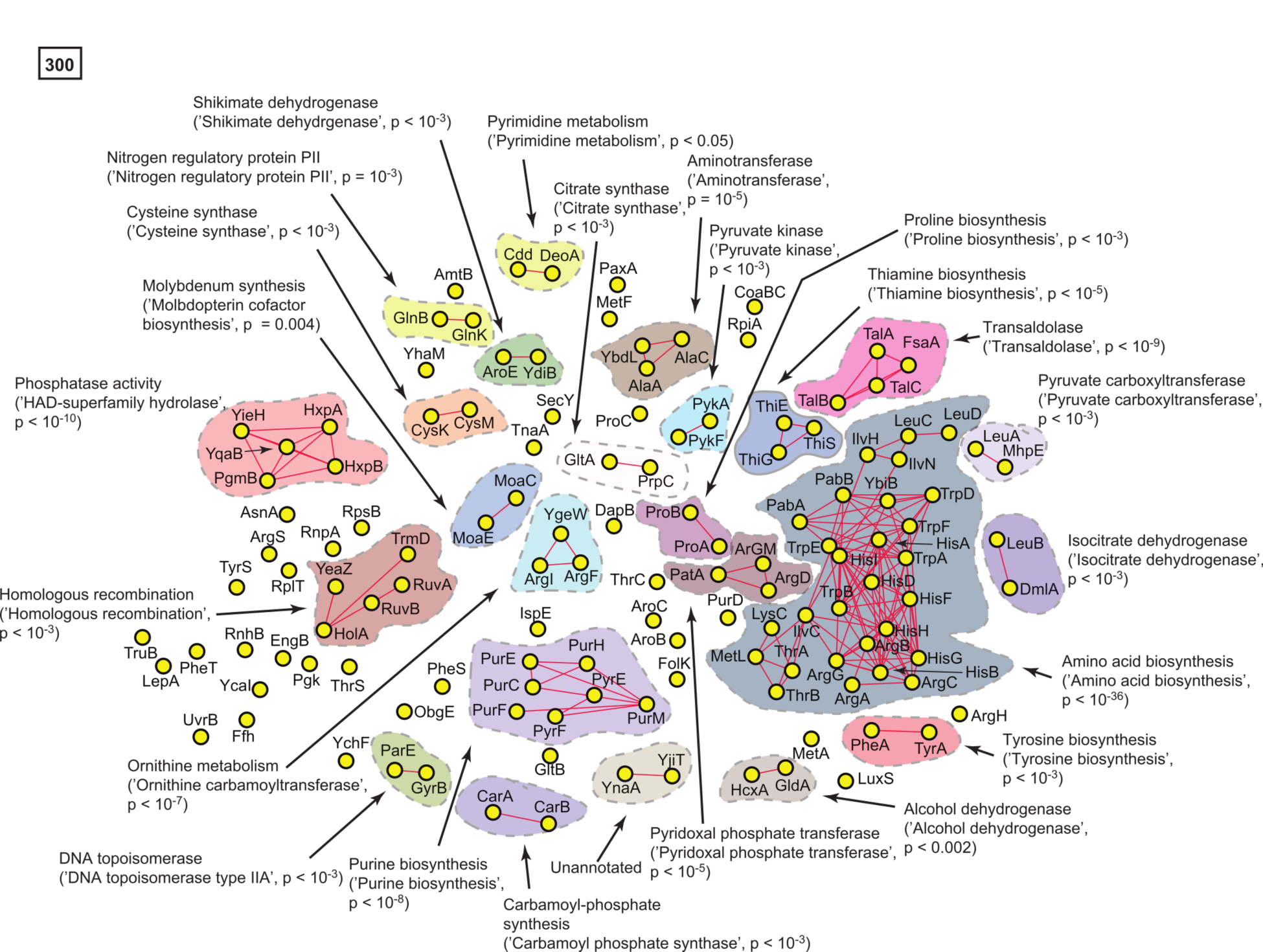
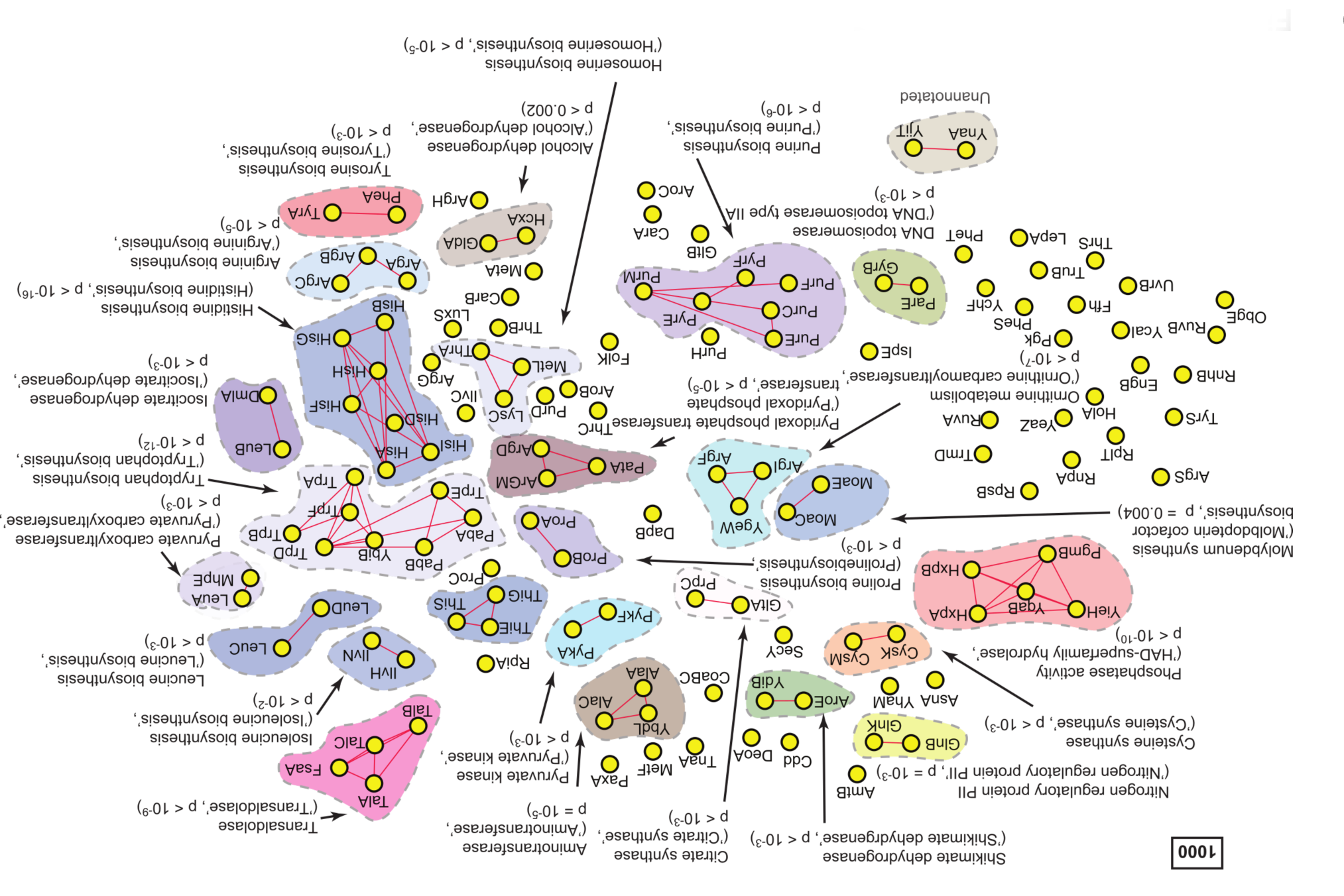
A statistically-derived hierarchical model of amino acid metabolism in *E. coli* K12 using HisG as a query protein. (**A**) 129 proteins in *E. coli* K12 shared significant spectral correlations with HisG across components SVD34 to SVD134. Statistical interaction networks were defined by thresholding spectral depth at 50, 150, 300, and 1000. Nodes (yellow circles) represent proteins and edges (red lines) represent statistical interactions. Shaded contours identify discrete subnetworks and are labeled with their assigned function based on interpretation of gene-set enrichment analysis (GSEA) and literature review. The most significantly enriched ontological term produced by GSEA and the associated p-value is shown in parentheses for each subnetwork (**Table S10**).

**Figure 4 – figure supplement 6.**
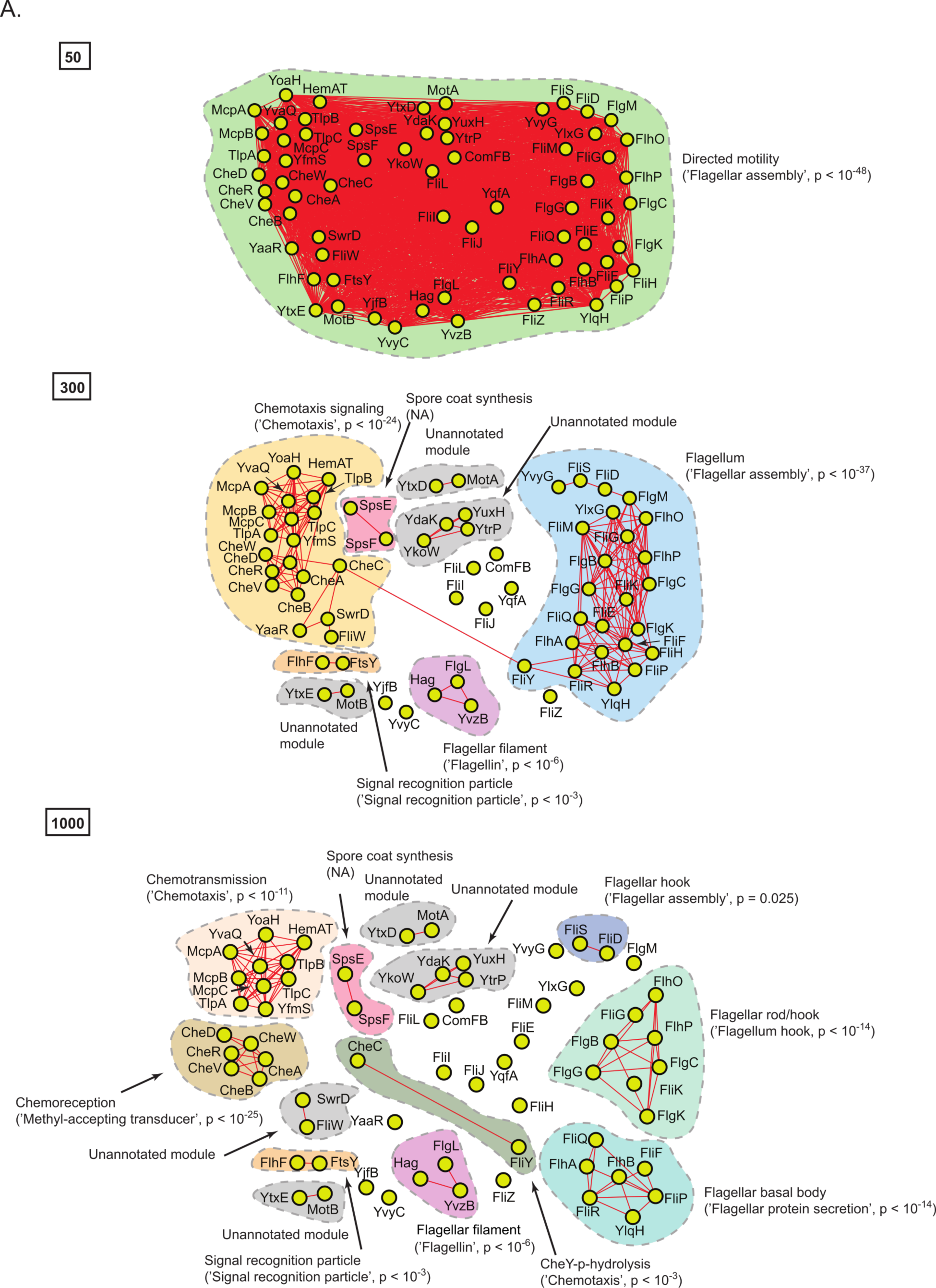
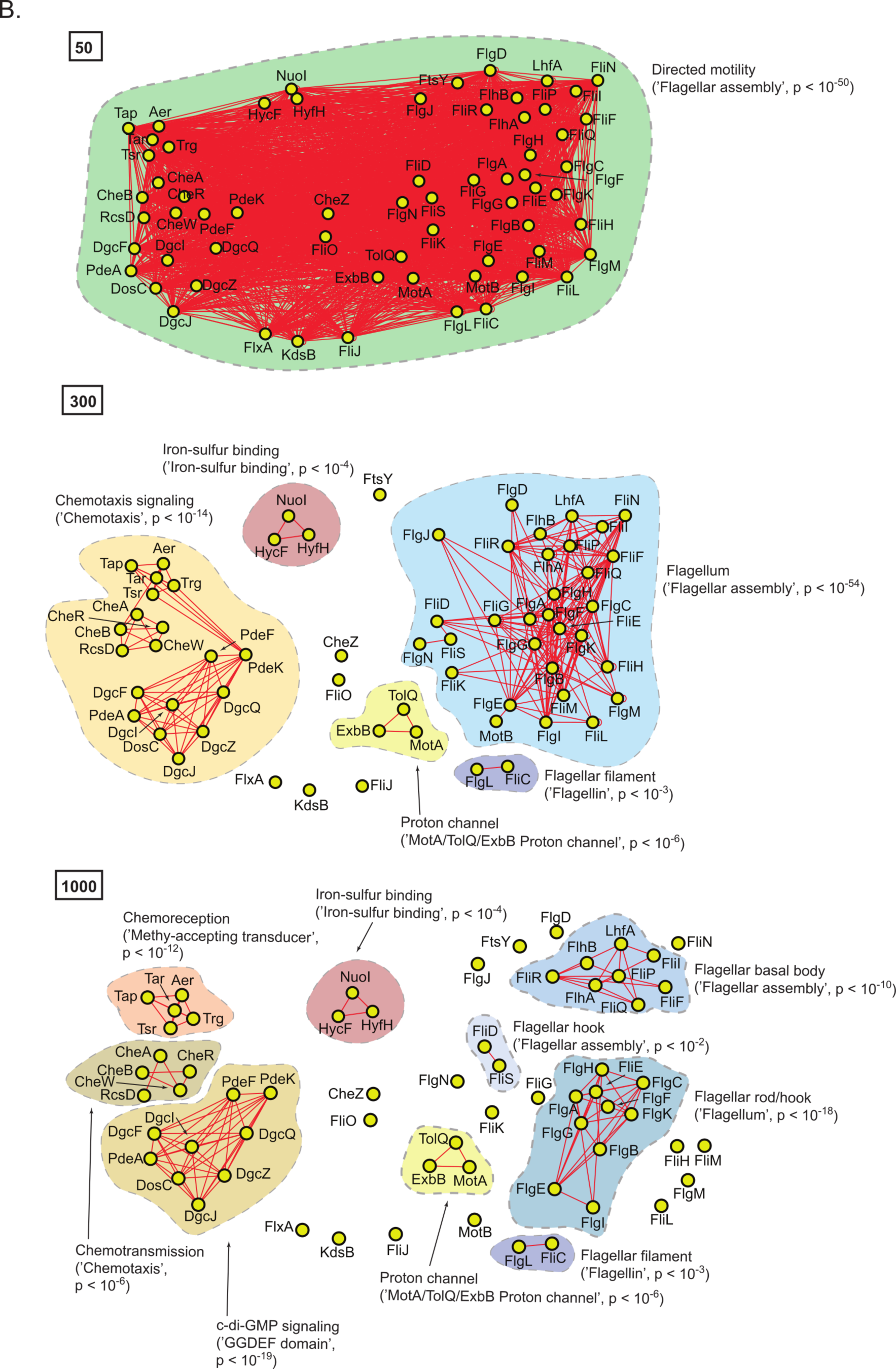
Statistically-derived hierarchical models of bacterial motility derived from serially thresholding spectral depth for correlations in *D^domain^*. (A) A model of motility in *B. subtilis* 168 using Hag as a query. (B) A model of motility in *E. coli* K12 using FliC as a query. Statistical interaction networks were defined by thresholding spectral depth at 50, 300, and 1000. Nodes (yellow circles) represent proteins and edges (red lines) represent statistical interactions. Shaded contours identify discrete subnetworks and are labeled with their assigned function based on interpretation of gene-set enrichment analysis (GSEA) and literature review. The most significantly enriched ontological term produced by GSEA and the associated p-value is shown in parentheses for each subnetwork (**Table S11, S12**).

**Figure 5 – figure supplement 1.**
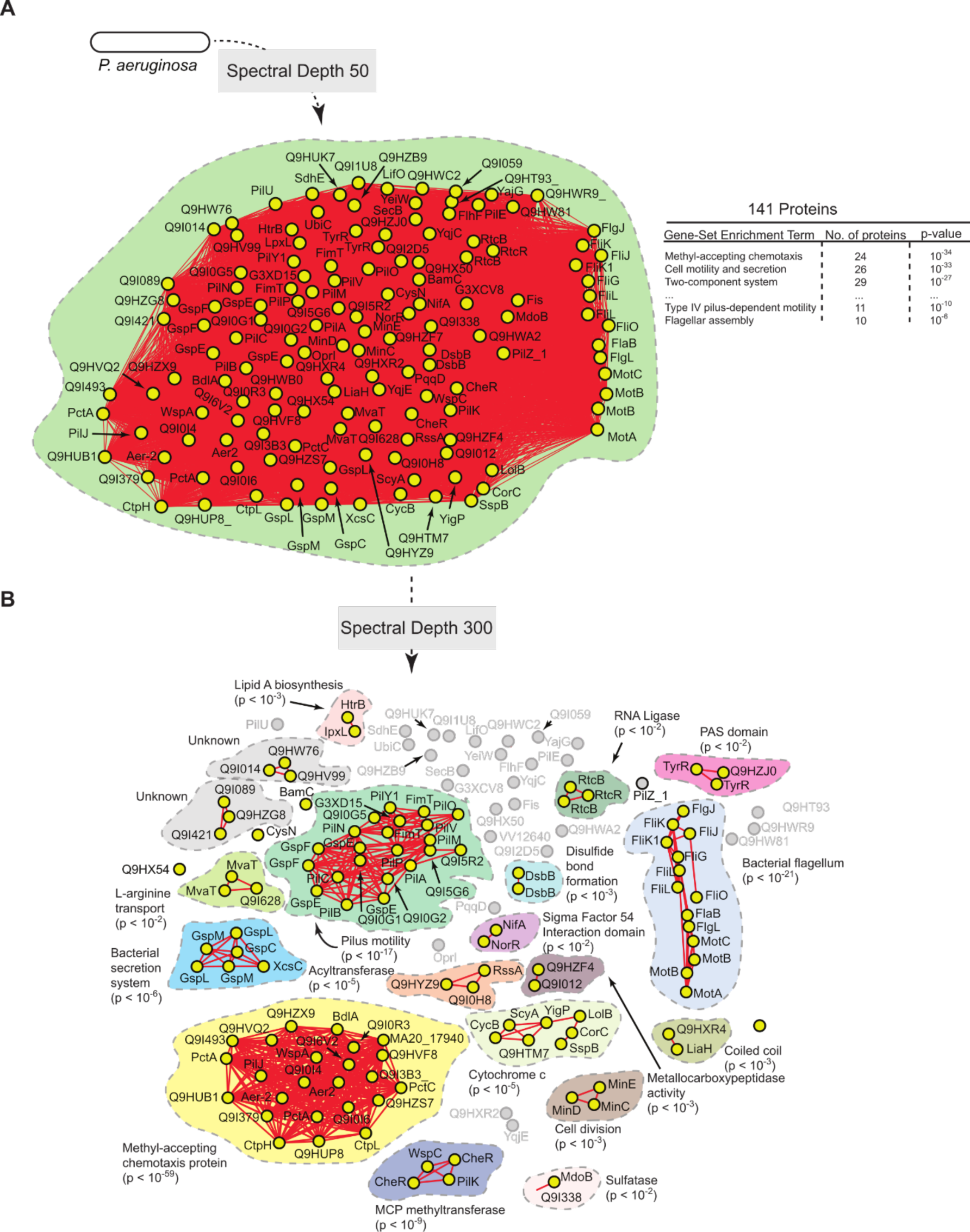
Statistically-derived hierarchical model of directed-motility in *P. aeruginosa* using PilA as a query. 140 proteins in *P. aeruginosa* shared significant spectral correlations with PilA across SVD34 to SVD134. (**A**) Statistical interaction network defined by thresholding spectral depth at 50. The inset illustrates significantly enriched terms resulting from gene-set enrichment analysis (GSEA) of the entire network. (**B**) Statistical interaction network defined by thresholding spectral depth at 300. Nodes (yellow circles) represent proteins and edges (red lines) represent statistical interactions. Shaded contours identify discrete subnetworks and are labeled in panel B with their assigned function based on interpretation of GSEA and literature review. The most significantly enriched ontological term produced by GSEA and the associated p-value is shown in parentheses for each subnetwork (**Table S13**).

## References

1. Barabasi AL, Zoltan O, (2004) **Network Biology: Understanding the cell’s functional organization** *Nat. Rev*. Gen. 5:101–113. https://doi.org/10.1038/nrg1272

2. Chuang HY, Hofree M, Ideker TA, (2010) **A decade of systems biology**. Annu. Rev. Cell Dev. Biol. 26:721–744. https://doi.org/10.1146/annurev-cellbio-100109-104122

3. Hartwell LH, Hopfield JJ, Leibler S, Murray AW, (1999) **From molecular to modular cell biology**. Nature 402:C47–C52. https://doi.org/10.1038/3501154

4. Costanzo M. et al, (2016) A global genetic interaction network maps a wiring diagram of cellular function. Science 353:aaf1420. https://doi.org/10.1126/science.aaf1420

5. Papin J, Reed J, Paulsson BO, (2004) **Hierarchical thinking in network biology: the unbiased modularization of biochemical networks**. Trends in Biochemical Sciences 29:641–647. https://doi.org/10.1016/j.tibs.2004.10.001

6. Ravasz E, (2009) **Detecting Hierarchical Modularity in Biological Networks**. Methods Mol Biol. 541:145–60. https://doi.org/10.1007/978-1-59745-243-4_7

7. Nurse P, (2008) **Life, logic and information**. Nature 454:424–426. https://doi.org/10.1038/454424a

8. Rajagopala S et al., (2014) **The binary protein-protein interaction landscape of Escherichia coli**. Nat. Biotechnol. 32:285–290. https://doi.org/10.1038/nbt.2831

9. Scheonrock A et al., (2017) **Evolution of protein-protein interaction networks in yeast**. PLoS One 12:e0171920. https://doi.org/10.1371/journal.pone.0171920

10. Hauser R et al., (2014) **A second-generation protein-protein interaction network of Helicobacter pylori**. Mol. Cell. Proteomics 13:1318–1329. https://doi.org/10.1074/mcp.O113.033571

11. Koo B et al., (2017) Construction and analysis of two genome-scale deletion libraries for *Bacillus subtilis*. Cell Syst. 4:291–305.e.7. https://doi.org/10.1016/j.cels.2016.12.013

12. Luck K et al., (2020) **A reference map of the human binary protein interactome**. Nature 580:402–408. https://doi.org/10.1038/s41586-020-2188-x

13. Eisen J, (1998) Phylgenomics: Improving functional predictions for uncharacterized genes by evolutionary analysis. Genome Research 8:163–167. https://doi.org/10.1101/gr.8.3.163

14. Pellegrini M et al., (1999) **Assigning protein functions by comparative genome analysis: Protein phylogenetic profiles**. Proc. Natl. Acad. Sci. 96:4285–4288. https://doi.org/10.1073/pnas.96.8.4285

15. Enright A, Iliopoulos I, Kyrpides N, Ouzounis C, (1999) **Protein interaction maps for complete genomes based on gene fusion events**. Nature 402:86–90. https://doi.org/10.1038/47056

16. Valencia A, Pazos F, (2002) **Computational methods for the prediction of protein interactions**. Curr. Opinion in Struct. Biol. 12: 368–373. https://doi.org/10.1016/s0959-440x(02)00333-0

17. Croce G, Gueudre T, Cuevas M, Keidel V, Figliuzzi M, Szurmant H, Weigt M, (2019) **A mult- scale coevolutionary approach to predict interactions between protein domains**. PLoS Comput. Biol. 15:e1006891. https://doi.org/10.1371/journal.pcbi.1006891

18. Cong Q, Anishchenko I, Ovchinnikov S, Baker D, (2019) **Protein interaction networks revealed by proteome coevolution**. Science 365:185–189. https://doi.org/10.1126/science.aaw6718

19. Green A, Elhabashy H, Brock K, Maddamsetti R, Kohlbacher O, Marks D, (2021) **Large- scale discovery of protein interactions at residue resolution using co-evolution calculated from genomic sequences**. Nat. Commun. 12:1396. https://doi.org/10.1093/bioinformatics/bty862

20. Szklarczyk D et al., (2018) STRING v11: protein-protein association networks with increased coverage, supporting functional discovery in genome-wide experimental datasets. Nucleic Acids Res. 47:D607–D613. https://doi.org/10.1093/nar/gky1131

21. Kuzmin E et al., (2018) **Systematic analysis of complex genetic interactions**. Science 360:eaao1729. https://doi.org/10.1126/science.aao1729

22. Kanehisa M, Goto S, (2000) **KEGG: Kyoto Encyclopedia of Genes and Genomes**. Nucleic Acids Res. 28:27–30. https://doi.org/10.1093/nar/28.1.27

23. Kanehisa M, (2019) **Toward understanding the origin and evolution of cellular organisms**. Protein Sci. 28:1947–1951. https://doi.org/10.1002/pro.3715

24. Kanehisa M, Furumichi M, Sato Y, Ishiguro-Watanabe M, Tanabe M, (2021) **KEGG: integrating viruses and cellular organisms**. Nucleic Acids Res. 49:D545–D551. https://doi.org/10.1093/nar/gkaa970.

25. Overbeek R, Fonstein M, D’Souza M, Pusch G, Maltsev N, (1999) **The use of gene clusters to infer functional coupling**. Proc. Nat’l. Acad. Sci. 96:2896–2901. https://doi.org/10.1073/pnas.96.6.2896

26. The UniProt Consortium, (2019) **UniProt: a worldwide hub of protein knowledge**. Nucleic Acids Res. 47:D506–515. https://doi.org/10.1093/nar/gky1049

27. Letunic I, Bork P, (2019) **Interactive Tree of Life (iTOL) v4: recent updates and new developments**. Nucleic Acids Res. 47:W256–W259. https://doi.org/10.1093/nar/gkz239

28. Huerta-Cepas J, Forslund K, Coelho LP, Szklarczyk D, Jensen L, Mering C, Bork P, (2017) **Fast Genome-Wide Functional Annotation through Orthology Assignment by eggNOG-Mapper**. Mol. Biol. Evol. 34:2115–2122. https://doi.org/10.1093/molbev/msx148

29. Huerta-Cepas J, et al. (2019) **eggNOG 5.0: a hierarchical, functionally and phylogenetically annotated orthology resource based on** 5090 **organisms and** 2502 **viruses**. Nucleic Acids Res. 47:D309–D314. https://doi.org/10.1093/nar/gky1085

30. Klema V, Laub A, (1980) **The singular value decomposition: Its computation and some applications**. IEEE Transactions on Automatic Control 25:164–176. https://doi.org/10.1109/TAC.1980.1102314

31. NCBI Resource Coordinators (2018) **Database resources of the National Center for Biotechnology Information**. Nucleic Acids Res. 46:D8–D13. https://doi.org/10.1093/nar/gkv1290

32. The Gene Ontology Consortium (2020) **The Gene Ontology resource: enriching a Gold mine**. Nucleic Acids Res. 49:D325–334. https://doi.org/10.1093/nar/gkaa1113

33. Kesler I, et al. (2016) **The EcoCyc database: reflecting new knowledge about Escherichia coli K-12**. Nucleic Acids Res. 45:D543–550. https://doi.org/10.1093/nar/gkw1003

34. Cong Q, Anishchenko I, Ovchinnikov S, Baker D, (2019) **Protein interaction networks revealed by proteome coevolution**. Science 365:185–189. https://doi.org/10.1126/science.aaw6718

35. Schafer, J., and Strimmer, K. (2005). **An empirical Bayes approach to inferring large- scale gene association networks**. Bioinformatics 6, 185–189. https://doi.org/10.1093/bioinformatics/bti062

36. Sul, J.H., Martin, L.S., and Eskin, E. (2018). **Population structure in genetic studies: confounding factors and mixed models**. PLoS. Genetics, http://doi.org/10.1371/journal.pgen.1007309

37. Nagy, L.G., Merenyi, Z., Hegedus, B, Balint, B. (2020). **Novel phylogenetic methods are needed for understanding gene function in the era of mega-scale genome sequencing**. Nucl. Acids. Res. 48, 2209–2219. https://doi.org/10.1093/nar/gkz1241

38. Babu, M. et al. (2014). Quantitative genome-wide genetic interaction screens reveal global epistatic relationships of protein complexes in *Escherichia coli*. PLoS Genetics 10, e1004120. https://doi.org/10.1371/journal.pgen.1004120

39. Babu, M. et al. (2018). **Global landscape of cell envelope protein complexes in Escherichia coli**. Nat. Biotechnol. 36, 103–112. https://doi.org/10.1038/nbt.4024

40. Hu, P. et al. (2009). **Global functional atlas of *Escherichia coli* encompassing previously uncharacterized proteins**. PLoS. Biol. 4, e100096. https://doi.org/10.1371/journal.pbio.1000096

41. Franceschini, A., Von Mering, C., Jensen, L.J. (2016). **SVD-phy: improved prediction of protein functional associations through singular value decomposition of phylogenetic profiles**. Bioinformatics 32, 1085–1087. https://doi.org/10.1093/bioinformatics/btv696

42. Mistry J, Finn R, (2007) **Pfam: a domain-centric method for analyzing proteins and proteomes**. Methods Mol. Biol. 396:43–58. https://doi.org/10.1007/978-1-59745-515-2_4

43. Huang, D.W., Sherma, B.T., Lempicki, R.A. (2009). **Bioinformatics enrichment tools: paths towards the comprehensive functional analysis of large gene lists**. Nucleic Acids Res. 37, 1–13. https://doi.org/10.1093/nar/gkn923

44. Huang, D.W., Sherma, B.T., Lempicki, R.A. (2009). **Systematic and integrative analysis of large gene lists using DAVID Bioinformatics Resources**. Nature Protoc. 4, 44–57. https://doi.org/10.1038/nprot.2008.211

45. Paul, K., Nieto, V., Carlquist, W.C., Blair, D.F., Harshey, R. (2010). **The c-di-GMP binding protein YcgR controls flagellar motor direction and speed to affect chemotaxis by a “Backstop Brake” mechanism**. Mol. Cell. 38, 128–139. https://doi.org/10.1016/j.molcel.2010.03.001

46. Walker, S.L., Redman, J.A., Elimelech, M. (2004). **Role of cell surface lipopolysaccharides in Escherichia coli K12 adhesion and transport**. Langmuir 18, 7736–7746 (2004). https://doi.org/10.1021/la049511f

47. Alm R, Mattick J (1995) Identification of a gene, pilV, required for type 4 fimbrial biogenesis in Pseudomonas aeruginosa, whose product possess a pre-pilin-like leader sequence. Mol. Microbiol. 16:485–496. https://doi.org/10.1111/j.1365-2958.1995.tb02413.x

48. Little A, et al. (2018) *Pseudomonas aeruginosa* AlgR phosphorylation status differentially regulates pyocyanin and pyoverdine production. mBio. 9:e02318–17. https://doi.org/10.1128/mBio.02318-17.

49. Kearns D, Robinson J, Shimkets L, (2001) **Pseudomonas aeruginosa exhibits directed twitching motility up phosphatidylethanolamine gradients**. J Bacteriol. 183:763–7. https://doi.org/10.1128/JB.183.2.763-767.2001

50. Rashid M, Kornberg A, (2000) Inorganic polyphosphate is needed for swimming, swarming, and twitching motilities of *Pseudomonas aeruginosa*. Proc. Natl. Acad. Sci. 97: 4885–4890. https://doi.org/10.1073/pnas.060030097

51. Wigner, E. P., (1967) **Random matrices in physics**. SIAM Rev., 9(1):1–23. https://doi.org/10.1137/1009001

52. Cover and Thomas (2006) Elements of information theory, 2nd edition. ISBN: 978-0-471- 24195-9

